# CompBio and MIRaS – A Multi-omic Analysis Platform Built on a Memory-Based Intelligence Engine

**DOI:** 10.64898/2025.12.11.693741

**Authors:** Ruteja A Barve, Chad E Storer, Wayne D Hoxsie, Curtis Marcum, Joshua F McMichael, Jared M Lalmansingh, Brittany K Smith, Mark R Johnson, Daniel J Kuster, Richard D Head

## Abstract

As molecular and cellular technologies have advanced, the need to analyze and interpret the resulting, often vast and multi-modal, data into actionable intelligence has become a rate-limiting factor in scientific advancement. While traditional knowledgebase-pathway tools and LLM-based AI tools both provide a degree of support in data interpretation, both exhibit limitations. Presented here is the CompBio multi-omic analysis platform built upon a novel memory-based intelligence engine, MIRaS. The system is not dependent on human-curated pathway knowledgebases, can perform statistical significance analysis, does not suffer from hallucination, and produces traceable results. Furthermore, substantial effort has been placed in the computer-human knowledge transfer components of the system to enable efficient interaction, interpretation, and learning. The system has undergone years of extensive testing, validation, and evolution in omics-based research with dozens of associated publications verifying its effectiveness. A systematic assessment of CompBio and a detailed description of the MIRaS method are provided.

## Introduction

Accurate interpretation of high-complexity omic and multi-omic data has become paramount for biomedical research in its quest to drive efficient and effective therapeutic intervention development and application in precision medicine. As molecular and cellular technologies have advanced, the quantity and plexity of these data have grown exponentially. To elucidate the connotations associated with these often vast numbers of molecular measurements and changes, researchers have traditionally relied on ontology or knowledgebase pathway and enrichment analysis tools (GSEA^1,2^, ToppGene^3–5^, EnrichR^6^). These systems aid in promoting human understanding of omics data through a reductionist approach. Typically transforming lists with hundreds or thousands of biological entities into subsequent lists of tens to hundreds of biological pathways and/or processes representing the putative function of these entities within the associated biological condition(s). Tools may differ in their algorithmic approach but the foundational reliance on curated pathway or process knowledgebases is common, and though they have demonstrated their value over the last two to three decades, they have well-understood limitations that can lead to potential discoveries being missed^7^. Additionally, as there is no single knowledgebase that captures the full breadth of all biological pathways and processes, pathway tools typically incorporate many resources for comprehensiveness (GO^8,9^, Reactome^10,11^, KEGG^12^), often leading to substantial redundancy significantly complicating the act of human interpretation. Finally, the pathways or processes reported can, in cases, be exceptionally general (e.g. *Cellular Process*) providing little if any useful information to the user. While newer computational tools have attempted to address some of these issues (e.g. Metascape’s ability to collapse redundant pathways)^13^, the differential between what is known and what is captured in the curated resources remains substantial.

Within the last few years, artificial intelligence-based omics analysis systems have begun to assert their position in the quest for biomedical knowledge and hypothesis generation^14–17^. Due to their ability to incorporate additional biological knowledge beyond the simple entity/list associations or even topological associations from canonical pathways, these platforms have substantially enhanced the interpretation processes. However, even with their extensive capabilities, these systems too have limitations. The most prominent of which is the inability to produce the statistical significance provided for pathways by traditional tools. Also, while these systems can provide exemplar resources to support a given finding or assertion, explicit tracing of the information used to derive a particular result is generally not possible. Combined with the difficulty of occasional hallucination, it is difficult to ascribe accurate confidence factors for AI-based omic or multi-omic analysis results involving LLMs or other models.

CompBio is designed to produce enhanced biological pathway and process analyses without dependence on pathway knowledgebases. To accomplish this, the analysis platform is built upon a novel episodic memory-based intelligence engine known as MIRaS. MIRaS encodes the information contained within knowledge sources such as PubMed abstracts and full-text articles as episodic memories into a framework capable of not only storing memories but also creating and comparing complex (semantic) memories for the purpose of inference and simple reasoning. Due to the ability of MIRaS to create randomized complex memories on demand, the system is capable of empirically-derived statistical significance on multiple levels of findings, including the full experiment level, and can trace all findings back to the exact source knowledge used in their formation. Moreover, the intermediary components designed to convert the system-derived knowledge into human interpretable information can not only produce the most likely biological pathways and processes represented by the input omics data but illustrate how those pathways and processes interconnect with each other through conversion of the initial knowledge graph into a 3-dimensional knowledge map representation for user exploration and learning.

Through its novel construction, CompBio addresses particular limitations of both traditional and more recent AI-based omics analysis tools including the lack of need for pathway knowledgebases, ability to store, compute on, and trace back to episodic memory, form complex, semantic memory with unlimited context, and generate p-values for findings. As successive and evolving versions of CompBio have been in use for nearly five years, dozens of high-impact publications have demonstrated the value of this construct with key findings in numerous areas of biomedical research^18–23^. While early versions of the CompBio and MIRaS technology were initially licensed for commercial application, system access is now freely available to all non-commercial users. Here is presented a systematic, results-centric assessment of CompBio and a detailed description of the MIRaS engine upon which it is built.

## Results

### A multi-omic analysis platform built on a memory-based intelligence engine

The CompBio platform is an application specifically designed to accelerate scientific research through multi-omic data analysis and interpretation. CompBio, in turn, is built upon a more generalizable AI platform described as a **M**emory-based **I**ntelligent **R**e**a**soning **S**ystem (**MIRaS**).

Within CompBio, a web-based client provides an integrated user environment that interfaces with the MIRaS server, as well as several post-processing features including advanced image search and chatbot tools to support user idea and hypothesis generation (**Figure 1**). MIRaS is a novel construct capable of ingesting and storing information as recallable memories from a highly efficient, multi-dimensional memory framework. Unlike model-based AI systems that train on information and/or data, MIRaS explicitly stores the knowledge it ingests, initially, in the form of episodic memories (**Figure 2a**). These episodic memories can, in turn, be used to form more complex memories (aggregations of logically or contextually related episodic memories) (**Figure 2b**). Subsequent normalization of complex memories generates semantic memories (**Figure 2c**). All episodic, complex, and semantic memories stored within the memory framework form the **Ge**neralized **M**emory **M**odel (**GeMM**). Instead of “predicting” a response to a user input based on prior training, the **M**emory-Based **I**nference and **R**easoning engine (**MIR**) will directly infer or “reason” a response, in near real time, through information-theoretic calculations, directly from the GeMM. Within this system, all forms of knowledge source-derived memories, can be recalled, further aggregated, normalized, and computed on to identify “signal”, or information relevant to user inquiries. The MIRaS that supports CompBio for multi-omic analysis ingests knowledge from PubMed abstracts and full-text articles. User inquiries, which typically consist of genes, proteins, or metabolites from biological data sets of interest, form MIRaS knowledge graphs (**Figure 3a**). Utilizing a knowledge conversion or transfer tool set, CompBio converts the MIRaS knowledge graph into an interactive knowledge map of inter-connected biological themes, higher-order constructs that represent biological pathways, processes, and cellular functions associated with the initial inquiry (**Figure 3b**). Due to the nature of the framework and computation thereon, all results are traceable to their explicit source knowledge. It is also possible to determine the statistical significance of an analysis and its related output derived from inquiries of the MIRaS. A further capability of the platform is the Assertion Engine (AE), an ML-based utility, which is capable of complex knowledge graph comparison for interpretive analysis. Unlike tools that perform simple subgraph detection, the AE assesses the degree and depth of contextual similarity for matching subgraph components. Moreover, this engine can compute an overall similarity between knowledge graphs and will also provide a level of statistical significance. As an exemplar application, AE can identify preserved biology between comparative data such as animal models and human disease. In effect, AE can make an “assertion” as to whether the model recapitulates aspects of disease and the degree of confidence that these aspects are non-random. Finally, given the generalizability of the framework and subsequent computation, the system can be, and has been, successfully applied to a multitude of applications extending well beyond multi-omic analysis including Electronic Medical Record (EMR) and FDA Adverse Event Reporting System (FAERS) data (not published here).

**Figure 1.**
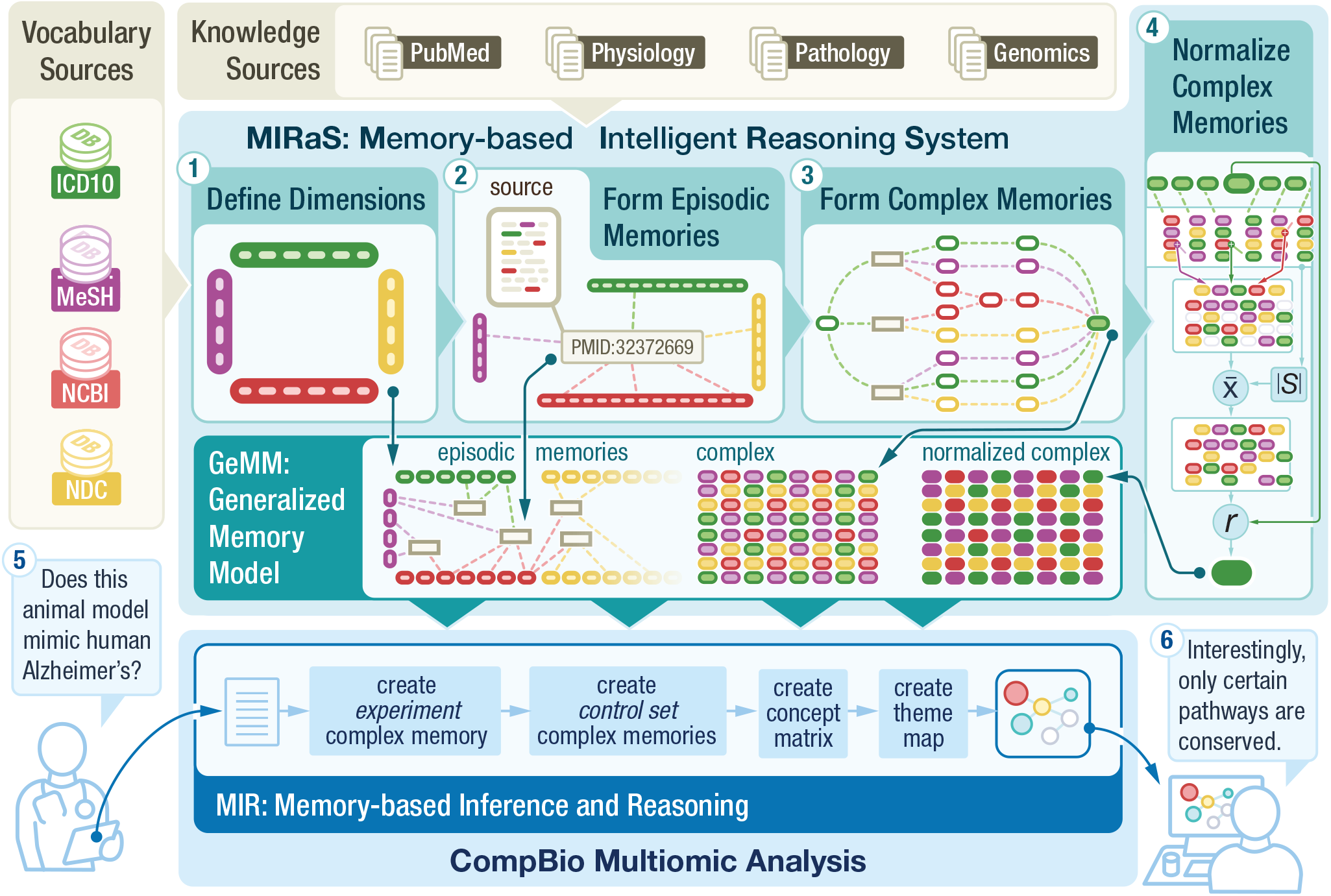
CompBio and MIRaS system overview. Overview of the three components: (1) framework definition, (2) source knowledge ingesting, and (3) complex memory formation used to build the MIRaS GeMM from source knowledge; and, the three components (4) memory-based reasoning, (5) system inquiry, and (6) knowledge transfer utilized to generate and return knowledge to the user.

**Figure 2.**
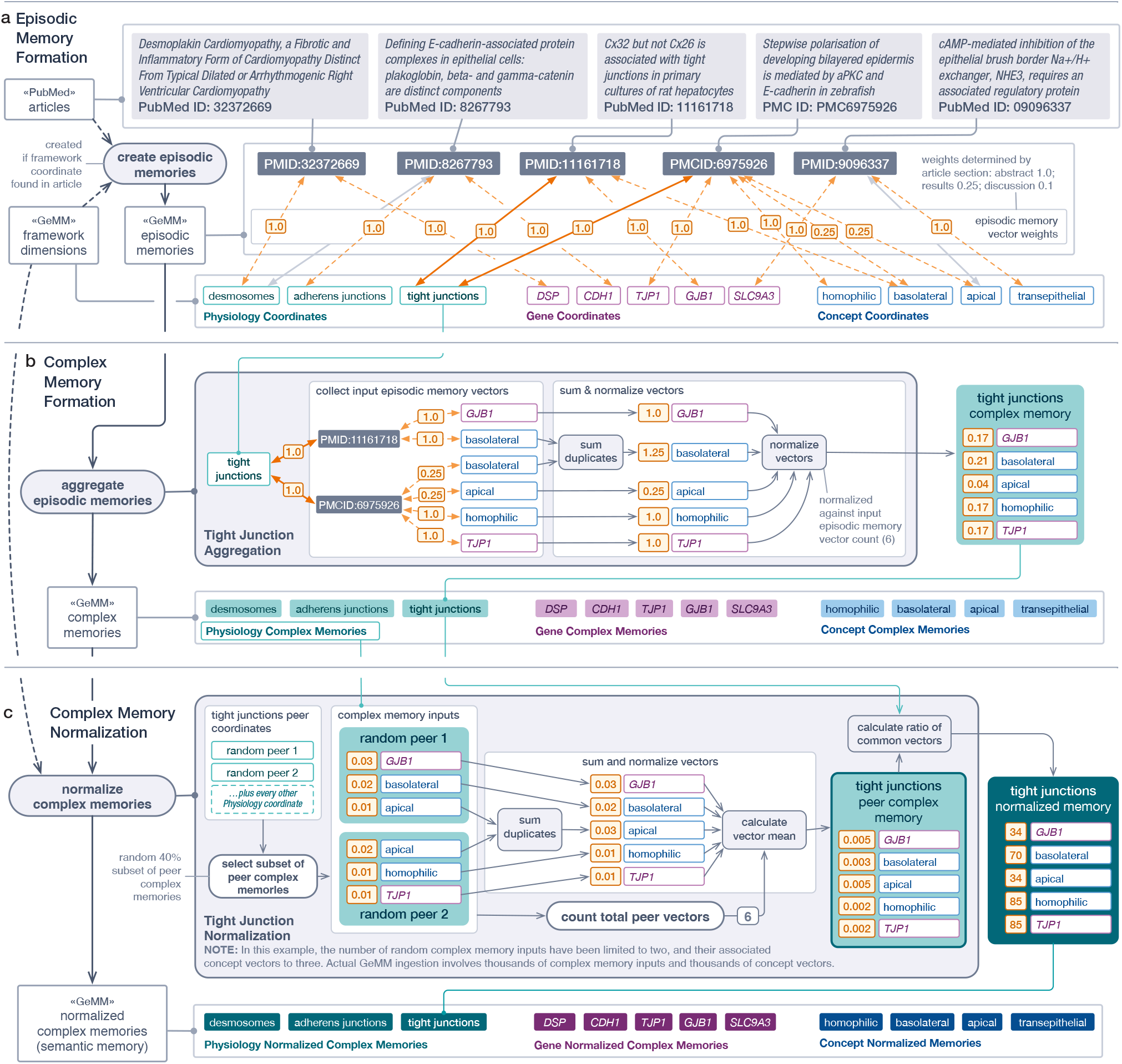
Conceptualization of CompBio GeMM generation. Once a knowledge framework is defined (**Supplemental Figure 1**), the generation of the MIRaS GeMM occurs through a three-step process. A highly-simplified conceptual illustration of these steps is provided here. (**a**.) An example of episodic memory formation based on a three-dimension framework (entities, physiology, and biological concepts). Each source of knowledge (PubMed abstract or article) is initially tokenized based on the vocabulary identified within the framework dimensions and then vectorized to create the stored episodic memory. (**b**.) Complex memories are formed for each dimensional coordinate within the knowledge framework through the aggregation of all episodic memories with a vector to that coordinate. In this specific illustration, all episodic memories containing the “tight_junctions” token are aggregated to form a complex memory for the tight_junctions coordinate within the physiology dimension. (**c**.) To identify potentially meaningful cross-coordinate associations within a complex memory (i.e. signal), the given complex memory for “tight junctions” is normalized by a randomly selected set of peer complex memories from the same dimension (physiology). The complex memory for “tight_junctions” is then normalized through a vector-by-vector comparison to the aggregated, then averaged, complex memory of the random peers. Thus, the GeMM is defined as the set of all episodic memories and all coordinate normalized complex memories constructed on the defined knowledge framework. For the CompBio GeMM, it is generated from 38+ million abstracts and 6.6+ million full-text articles. This process takes less than one day of real time on a single compute server conforming to the system requirements provided in the methods.

**Figure 3.**
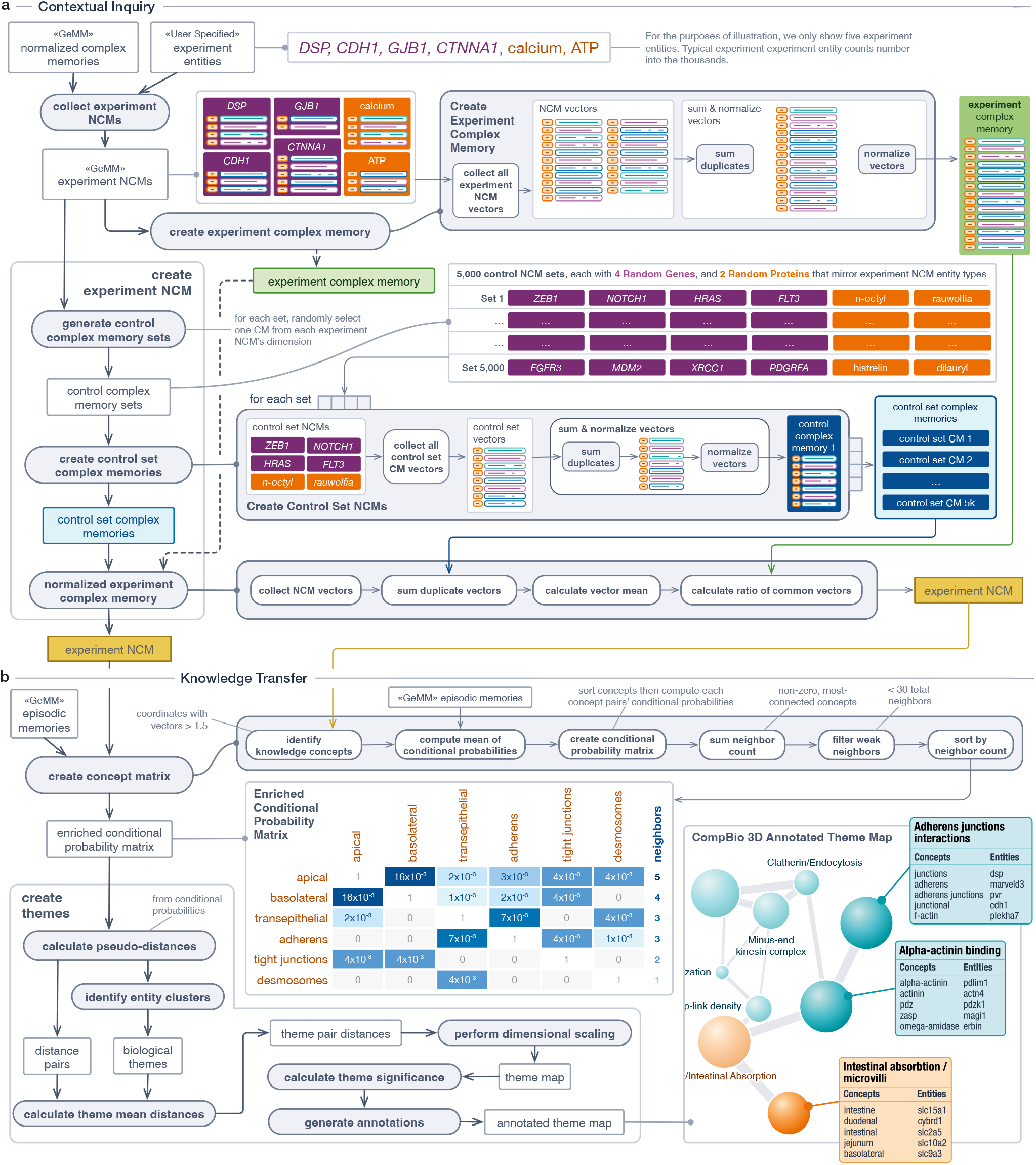
System inquiry and knowledge-graph generation for CompBio. GeMM generation occurs once for a given source knowledge set. Only when new source knowledge is added is the process reinitiated. By design, the GeMM stays resident in system memory to provide “memory-as-a-service” to the MIR component of MIRaS. (**a**.) A high-level overview of the system workflow to process inquiries from CompBio users. Leveraging the same, flexible process as GeMM creation, MIR generates an experiment-level complex memory. The entities and entity types are immediately identified to create peer memories from randomized entities of the same number and type (i.e. peers), as the user query. For example, if the user inquiry consists of 400 genes and 100 metabolites, randomized comparators will contain identical numbers of genes and metabolites that are non-overlapping with the input set(s). The inquiry complex memory is then normalized to the mean complex memory of the 5,000 random comparators to identify inquiry-specific signal information (**b**.) For CompBio, the normalized complex memory (NCM) of the experiment represents the contextual knowledge generated by MIR based on the user inquiry that, in its raw form, would be uninterpretable by humans. The knowledge transfer tool set converts the MIR information, initially represented as a knowledge graph, into an explorable 3 - dimensional knowledge map of labeled biological themes which are interconnected based on their biological relationships. Through an approach described in the methods, p-values are computed at both the overall experiment and individual theme levels.

To establish the utility and accuracy of CompBio and, by extension, MIRaS, three levels of assessment were conducted. First, as MIRaS has no knowledge of entity-pathway associations (genes, proteins, metabolites) stored in pathway knowledgebases, a series of positive control assessments were performed to demonstrate the ability to correctly identify established biology. Second, a set of comparisons with traditional pathway tools were undertaken on published data to demonstrate how CompBio compares to designed-for-purpose tools that are statistical-significance capable. Finally, as versions of CompBio have been in use for some time, real-world evidence as to the impact of the platform from prospective validation of findings will be presented to demonstrate the systems abilities to aid in reliable hypothesis generation. These results are provided in the context of traditional pathway tools and a more modern LLM based platform.

### Positive control assessment – Hallmark pathway recognition

Because the MIRaS engine does not rely on curated entity-pathway resources, we benchmarked recovery of established pathways from their gene lists using MSigDB Hallmark collection^24^, which comprises consensus, expert-curated gene sets aggregated from multiple founder sets. CompBio results were evaluated as described in Methods, with 46 of 50 gene sets correctly identified (**Figure 4a, Supporting Data STable 1**).

**Figure 4.**
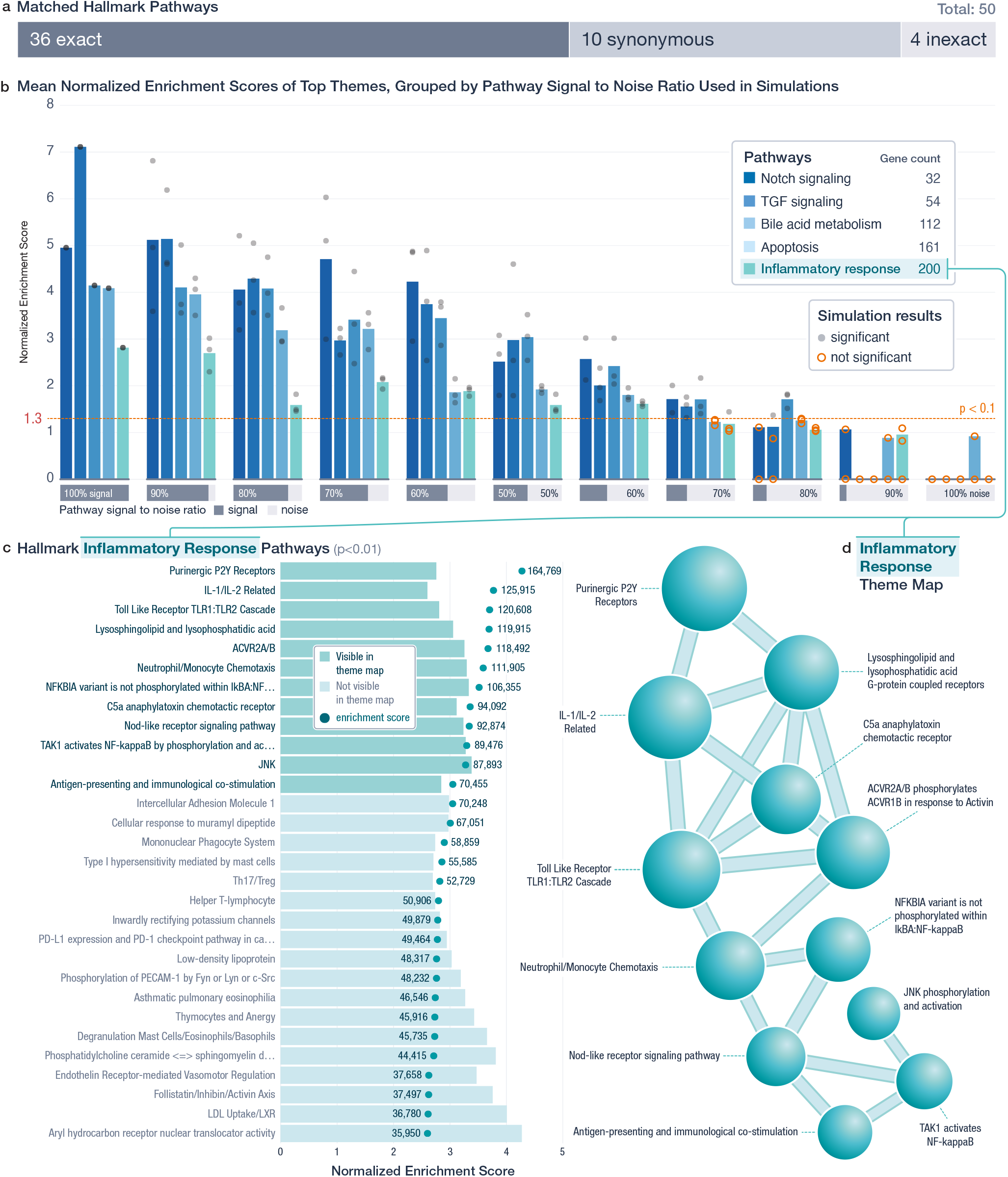
Positive-control assessment – Hallmark pathway gene sets. The Hallmark gene sets were utilized as a positive-control assessment to determine if CompBio, which has no knowledge of the entity-pathway associations from pathway knowledgebase, can reproduce the known biology associated with these sets. (**a**.) Expert review results of the individual CompBio maps generated for each of the 50 hallmark pathway genes sets. 46 of the 50 pathways produced an exact or synonymous result at the theme level, concept level or both. The four sets of genes without an exact or synonymous match are, notably, up- and down-regulated gene lists from two experimental conditions that do not represent specific pathways or processes. (**b**.) To assess signal-to-noise tolerance, five hallmark pathways were subjected to progressive noise introduction with concordant signal reduction (0–100% signal, x-axis), in which pathway genes were randomly replaced with genes not in the Hallmark set(s). The bar graph illustrates the mean normalized enrichment scores (NES) of CompBio themes matching the corresponding Hallmark pathway, while the associate circles indicate the p-value for that theme. Each signal level includes three randomized trials per pathway. Grey circles denote p-values < 0.1 while orange circles represent non-significant matches; a missing bar indicates no thematic match to the Hallmark pathway. (**c**.) Detailed CompBio analysis and results of the Inflammatory Response gene set (top 30 significant themes), ranked by enrichment score. Filled circles indicate enrichment scores by which themes are initially ranked for sizing and p-value calculation; darker bars denote themes also present in the adjacent knowledge map view. (**d**.) CompBio knowledge map view of the Hallmark *Inflammatory Response* gene list that illustrates both the spatial positioning of themes (spheres) and their interconnections (edges), based on their “biological closeness” and shared entities as determined by MIRaS.

### Positive control assessment – signal/noise tolerance, statistical significance

To assess robustness, we replaced fixed fractions of genes in selected Hallmark sets with randomly sampled non-pathway genes, while maintaining consistent overall list size. The noise fraction was increased in 10% increments (0-100%), each in triplicate. Across five pathways, the normalized enrichment score (NES) declined nearly linearly with noise (**Figure 4b, Supporting Data STable 2**). Gene sets with > 20% signal typically maintained significance (p≤0.1; NES≥1.3), beyond this most replicates fell below threshold, and at 100% noise no significant signal was detected, indicating that performance declines with increasing signal corruption.

CompBio’s ability to recover peripheral, contextually related processes, and interrelationships, is shown by the ranked top-30 theme list (enrichment and NES; **Figure 4c**) and the corresponding knowledge map illustrating inter-theme relatedness (**Figure 4d, Supplemental Figure 2**,**Supporting Data STable 3**).

### Positive control assessment – biological data with known mechanisms

We benchmarked CompBio on publicly available transcriptomic data from primary human cells treated with LOPAC1280 compounds^25^ with established mechanisms of action (MoA), aiming to recover tool compound MoA from expression signatures and to reveal drug-class mechanistic overlap. CompBio analysis of signatures from dermal fibroblasts and aortic smooth muscle cells treated with brefeldin A (28.5nM and 900nM) recovered ER/Golgi trafficking, unfolded-protein response, and cilium/centriole theme consistent with prior reports^26,27^ (**Figure 5a, Supporting Data STable 4**). Contextual similarity was significant across all comparisons (p≤0.001) demonstrating conservation of compound-driven mechanistic signal. The strongest similarities were observed within cell types, as expected (**Figure 5b**). A simplified AssertionEngine overlap map (**Figure 5c, Supplemental Figure 3**) shows a selection of conserved and inter-related concepts associated with the expected biological MoA for brefeldin A, observed across both cell types.

**Figure 5.**
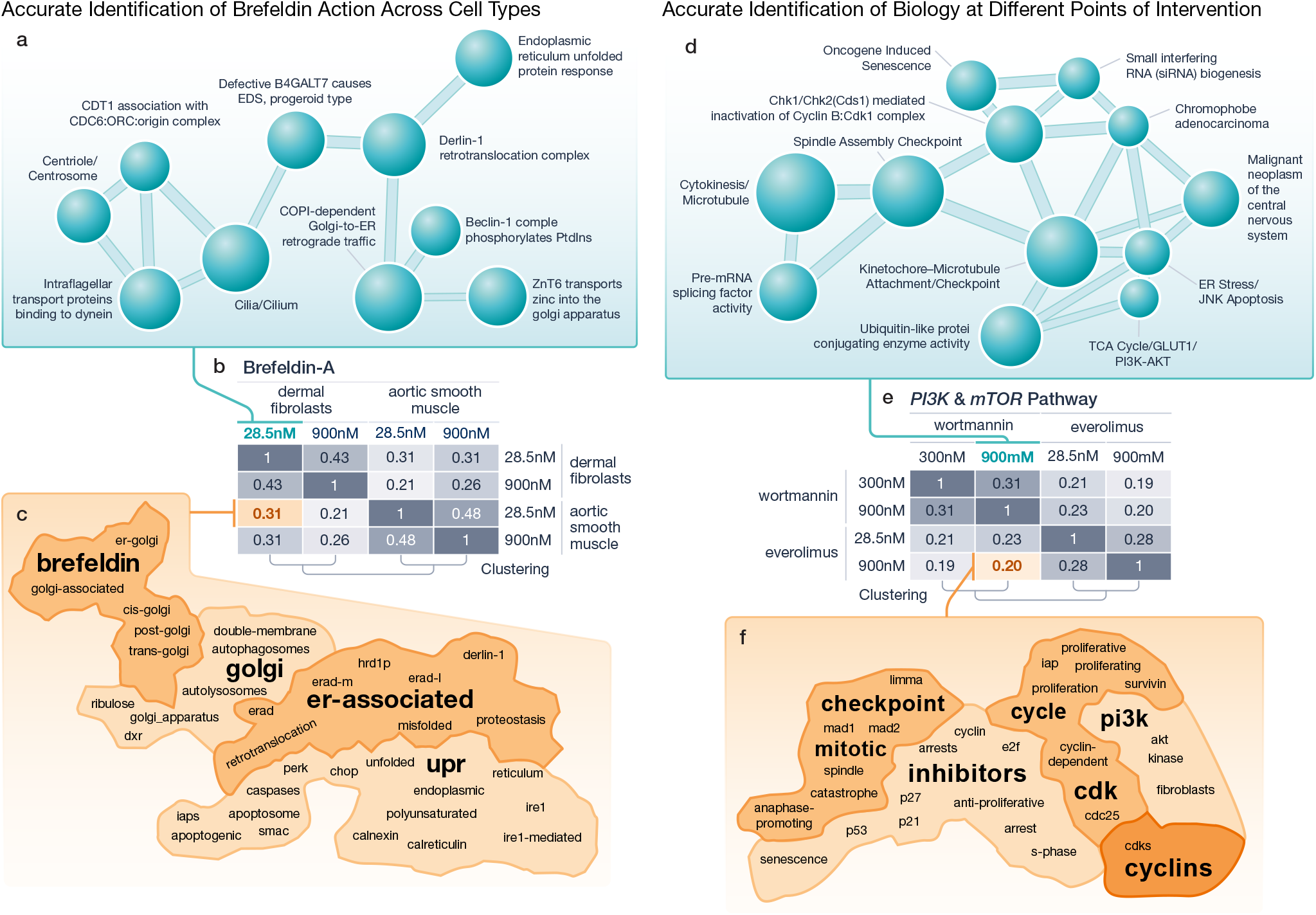
Positive-control assessment – biological data with known mechanisms of action. To further test the ability of CompBio to identify known biology, but on an experimental scale, RNA-Seq data from compounds with known mechanisms of action were rigorously assessed. Panels **a-c** provide results for brefeldin-A, a known inducer of ER stress and unfolded-protein response, across two cell types (dermal fibroblasts and aortic smooth muscle cells). Panels **d-e** compare compounds with targets in a common pathway, wortmannin (PI3K) and everolimus (mTOR), within a given cell type (dermal fibroblasts). (**a**.) Core themes in the CompBio knowledge map generated from upregulated genes in dermal fibroblasts treated with 28.5nM brefeldin-A, highlight ER–Golgi transport and unfolded protein response. (**b**.) AssertionEngine results (grid) comparing CompBio knowledge maps from fibroblasts and aortic smooth muscle cells treated with brefeldin-A at two concentrations (28.5nM or 900nM). Darker shades of grey in the matrix indicate stronger similarity between two maps. The strongest similarity occurs between maps of a given cell type regardless of concentration, while for comparisons with other cell types, there is a weaker but still significant similarity. Additionally, all comparisons performed were significant (p < 0.001). These results suggest that, at the concentrations assessed, a greater degree of biological response variation occurs due to cell type rather than concentration. (**c**.) AssertionEngine subgraph showing shared concepts between fibroblast maps from 28.5nM and 900nM brefeldin-A (orange highlighted cell, panel **b**). This indicates that concepts associated with the known mechanism of action were strongly and contextually preserved across CompBio knowledge maps. MoA (Mechanism of action) related terms are called out with larger font for readability. (**d**.) Core themes in the CompBio knowledge map of upregulated genes in fibroblasts treated with wortmannin (900nM) with PI3K association. The theme in the lower right section is PI3K-specific. (**e**.) AssertionEngine results comparing fibroblast maps from wortmannin and everolimus (28.5nM or 900nM). Darker shades of grey in the matrix indicate the strength of similarity between compared knowledge maps (p < 0.001 for all comparisons). In this case, the strongest overlaps were, as expected, observed with the drug and its target. (**f**.) AssertionEngine subgraph illustrating contextually conserved concepts in overlap, between 900nM wortmannin and 900nM everolimus. Interestingly, though everolimus inhibits just downstream, the PI3K-AKT component of the pathway was strongly preserved.

To assess drug-class mechanistic overlap, we used transcriptomic data from primary dermal fibroblasts treated with wortmannin (PI3K inhibitor, 300nM and 900nM) and everolimus (mTOR inhibitor, 28.5nM and 900nM). CompBio identified the expected PI3K/AKT/mTOR-axis themes related to PI3K-AKT inhibition^28^, PIP3/ AKT activity^29^, and downstream effects on proliferation and survival^30^ (**Figure 5d, Supporting Data STable 5**). Contextual similarity was significant across all comparisons (p≤0.001) with the greatest magnitudes occurring with the same drug across doses (**Figure 5e**). A simplified AssertionEngine overlap map (**Figure 5f, Supplemental Figure 4)** demonstrates conserved selected concepts consistent with shared PI3K/AKT/mTOR-axis inhibition across both drug treatments.

### Comparative assessment – traditional pathway tool evaluation

A comparative assessment for pathway mapping efficiency was performed with CompBio and three widely-used traditional pathway tools (Metascape, ToppGene, and GSEA) using three published datasets^2,31,32^. Efficiency was defined as the sensitivity in the number of input genes mapped to at least one pathway or theme and specificity (data reduction efficiency) is the number of pathways or themes required to explain the data set. All datasets were analyzed with the four tools using recommended parameters as described in the methods section with the resulting efficiency graphs (**Figure 6a-c, Supporting Data STable 6-13**). While each of the four tools had overlapping and unique resulting pathways, in all three cases CompBio outperformed the other tools in combined efficiency. Manual inspection of the resulting analyses, **Figure 6d-f**, illustrate that CompBio, even in the context of its efficient mapping, was able to identify all, or the majority of, the original findings in all three papers.

**Figure 6.**
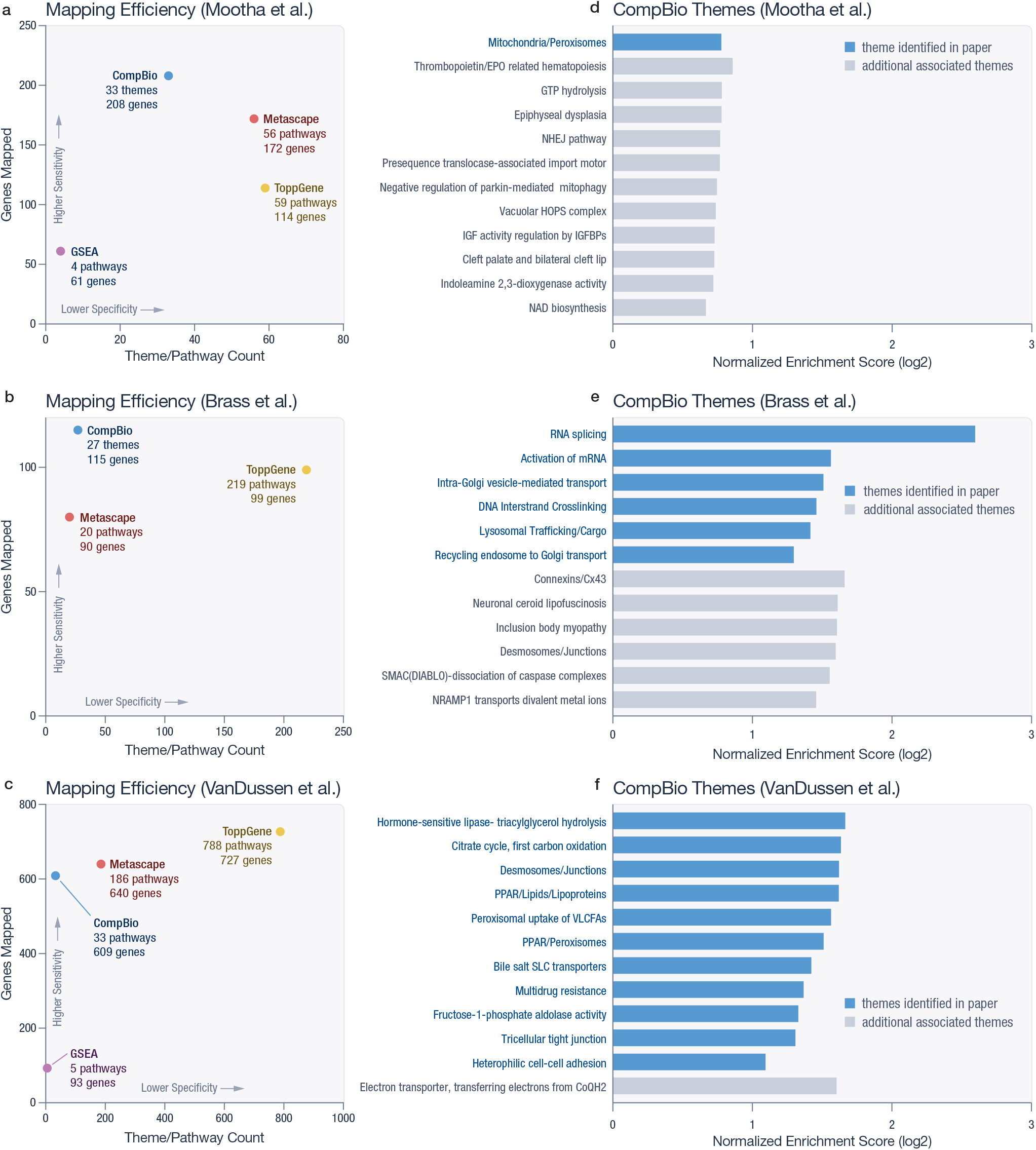
Comparative assessment – comparison with traditional pathway tools on published data. As an assessment for the key outcome, mapping efficiency, CompBio was tested with three traditional pathway tools (GSEA, ToppGene, and Metascape) to evaluate mapping efficiency. Mapping efficiency is defined as the ability to assign putative function to the most entities possible (sensitivity), with an optimal set of biological function descriptors, such as pathways and processes (specificity). The mapping efficiency was estimated utilizing three published data sets, including those used in the original publications of GSEA (Mootha et al.) and Metascape (Brass et al.) (**a-c**.) Mapping efficiencies for each tool on the respective published data, with the upper left quadrant indicating high efficiency. For the Mootha et al. dataset **(a.)** evaluation, CompBio mapped 208 of the 296 input genes into 33 themes outperforming Metascape (172 genes, 56 pathways), ToppGene (114 genes, 59 pathways), in both sensitivity and specificity. While GSEA (61 genes, 4 pathways) performed slightly higher in specificity, it had very low sensitivity, mapping only 61 of the original 296 genes. The Brass et al. data set demonstrated similar results (**b**.) CompBio mapped 115 out of 122 genes into 27 themes, compared with Metascape (80 genes; 20 pathways), and ToppGene (99 genes; 219 pathways). In this case, ToppGene actually demonstrated data inflation as it produced over twice as many significant pathways as input genes. This dataset could not be analyzed with GSEA because it lacked a ranking metric such as fold change. In the Vandussen et al. dataset (**c**.), CompBio mapped 609 genes out of the 1056 input genes with 33 themes, compared with Metascape (640 genes; 186 pathways), ToppGene (727 genes; 788 pathways), and GSEA (93 genes; 5 pathways). In this single case, Metascape and ToppGene mapped 3% and 9% more of the input gene list, respectively, than CompBio but required ~5x and ~24x as many pathways to do so. In all three data sets, CompBio consistently optimizes efficiency on both axes (sensitivity and specificity) while traditional tools typically optimize on only one axis, though Metascape performs closely to CompBio when using the collapse function in two cases. (NOTE: due to the nature of the input data set, GSEA could not be used to evaluate the Brass et al. data set.) (**d-f**) The top themes identified by CompBio for each corresponding dataset demonstrate that CompBio is able to identify all, or most, of the reported findings from the original publications (blue bars) with some additional findings (grey bars). Furthermore, the themes identified are clearly specific biological processes, not large, nondescript pathways or processes that simply capture large numbers of genes.

### Comparative assessment – an exemplar detailed pathway assessment

To provide an illustration of the limitation of knowledgebase-restricted tools in depth of knowledge recovery, all tools were cross-compared for gene recovery/association with an exemplar pathway. The adherens junction pathway, a key finding in the Vandussen et al. dataset, was identified by three of the four tools (not detected with GSEA). Gene membership assigned to the adherens junction concept (Heterophilic cell-cell adhesion theme) or pathway was compared across these three tools (**Figure 7, Supporting Data STable 14**). While only 5 genes overlapped between all three tools (**Figure 7b, 7d**), CompBio identified 10 of 11 Metascape genes and 19 of 24 ToppGene genes. Within the genes identified by CompBio, multiple well-characterized adherens junction associated genes such as *MYZAP*^*33,34*^, *TJP3*^*35*^, *DSG2*^*36*^ were not identified by the other tools.

**Figure 7.**
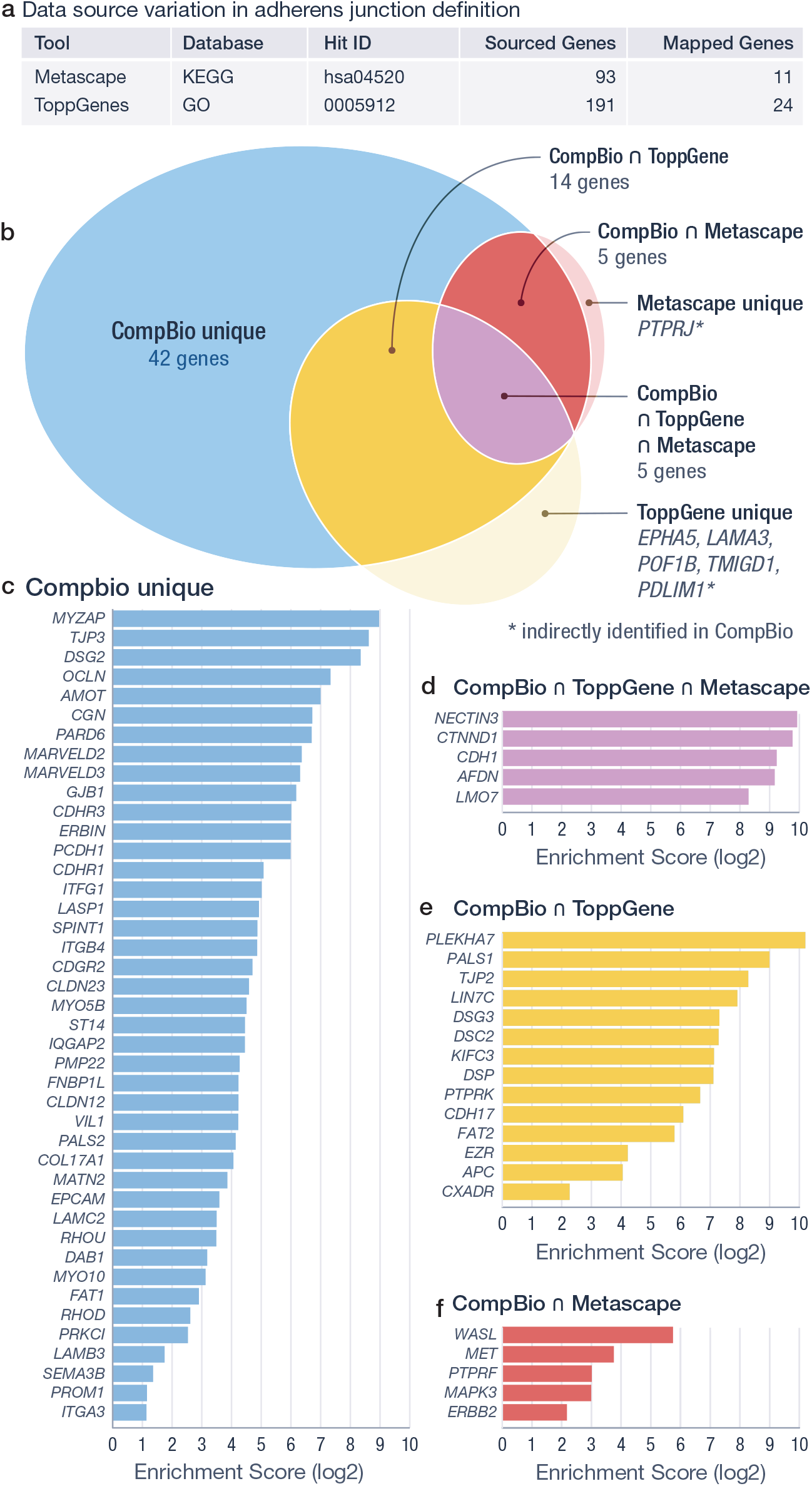
Comparative assessment – comparison with traditional pathway tools in a detailed pathway assessment. To illustrate the similarities and differences in how biological pathways and processes are identified between traditional curated knowledgebase tools and CompBio’s intelligence engine, the adherens junction pathway identified by Metascape, ToppGene, and CompBio (VanDussen et al.) was assessed in detail. (**a**.) Details of the specific pathway identification of adherens junction in Metascape and ToppGene. Though both have substantial overlaps in the underlying pathway knowledgebases used, Metascape identified the pathway from KEGG (hsa04520, −log(p) = 3.81), while ToppGene identified it from GO GO:0005912, FDR q-value [B&H] = 2.03 × 10^−4^). As a result, the source genes associated with pathway have substantial differences, as do the mapped genes. While GO and KEGG have 191 and 93 source genes for the pathway, respectively, only 21 overlap. Within the CompBio analysis, “adherens junction” was identified in the first theme (p = 1.84 × 10^−3^, NES = 2.159, concept enrichment score = 9225.25). (**b**.) Venn diagram for the genes identified by each tool showing uniqueness as well as overlapping genes. Using the VanDussen et al. data for the adherens junction pathway, the three tools yielded significant differences in the number of matching genes: Metascape (11), ToppGene (24), and CompBio (66). CompBio associated the majority of both the Metascape (10/11) and ToppGene (19/24) identified genes, while the overlap between Metascape and ToppGene was more modest. (**c**.) 42 genes, and their associated enrichment scores, uniquely identified as associated with the specific concept of adherens junction. Enrichment scores indicate the degree of association of each gene with the pathway. (**d-f**) Genes shared in the different tool intersections with their associated CompBio enrichment scores for adherens junction.

### Prospective assessment – laboratory validation of findings

As the CompBio platform has been in use with published findings since 2020^18,37,38^, it is possible to assess potential impact through reported findings that were validated in the laboratory prospectively. Three published examples of initial experimental findings that received subsequent laboratory corroboration or validation are presented here. Furthermore, the ability of other traditional tools to identify the comparable findings to CompBio, or not, with those specific data sets is also evaluated. To that end, Metascape and ToppGene were taken forward in this final assessment as they provided the strongest comparative results from traditional tools in the previous assessment. GeneAgent^14^, a new LLM-based tool for functional annotation of genes, was also tested.

### Assessment 1 – Sox9 intervertebral disc program (Tsingas et al.)

Tsingas et al. studied the effects of Sox9 deletion in the intervertebral disc of developing mice^39^. CompBio analysis of RNA-Seq data from disc compartments at 7 days illuminated substantial ECM changes in both studied compartments that preceded any visible histological effect.

Subsequent histological analysis of matching samples collected at 2 months and 7 months post Sox9 deletion revealed these early molecular changes did result in later phenotypic differences, corroborating the earlier CompBio results. Reanalysis of the 7-day dataset with Metascape identified a modestly related pathway “cartilage development” (GO:0051216, −log(p)=13.31) as the top hit among 24 collapsed pathways (**Supporting Data STable 16-1**). If the collapsing feature was not utilized (not recommended by the authors), a closely related pathway, “extracellular matrix structural constituent conferring tensile strength” (−log(p)=7.84) was identified. ToppGene explicitly identified “ECM structural constituent conferring tensile strength” (GO:0030020; FDR q-value [B&H] = 1.431×10^−6^) and “ECM” (GO:0005201, FDR q-value [B&H] = 1.43 × 10^−6^) (**Supporting Data STable 15**,**16-2**). GeneAgent also identified the process “Cartilage Development” (**Supporting Data STable 19-1**) indicating genes involved in regulation of cartilage development and ECM organization. In this case, ToppGene and GeneAgent accurately replicated the early finding, while Metascape’s collapsed results were related but only became explicit when collapsing was not employed.

### Assessment 2 – TH9 cells in LTBI and Mtb vaccination (Silver et al.)

In 2023, Silver et al. utilized CompBio to identify an unanticipated IL-9 expressing TH9-cell signature in latent tuberculosis as well as specific routes of BCG vaccination^40^. In addition to the original publications protein-based IL-9 confirmations, a subsequent publication demonstrated that the IL-9 producing TH9 cells are protective in an Mtb infection environment^41^. Metascape reanalysis of the Silver et al. data failed to identify any IL-9 or TH9 signatures within the data (**Supporting Data STable 17-1**,**17-2**). ToppGene reanalysis did not identify the TH9 cell type, but “IL-9 receptor binding” was present as pathway 34 of 58 by p-value (p=1.60 × 10^−3^; FDR q-value [B&H] = 3.95 × 10^−2^). GeneAgent identified “lymphocyte mediated immunity” in the final summarization step and IL-9 was only mentioned in the context general cytokine signaling and immune modulation with no mention of TH9 cells (**Supporting Data STable 19-2**). Thus, one traditional pathway tool would have missed the finding entirely while the other traditional pathway tool and LLM method would have required a substantial interpretive leap to reach the published CompBio finding.

### Assessment 3 – NAD+ synthesis and bile acid in EED (Malique & Sun et al.)

In 2024, Malique & Sun et al. demonstrated with CompBio that NAD+ synthesis and bile acid were common key pathway features down regulated in the ileum of both children with EED and a potentially translatable low-protein diet mouse model^42^. A prospective intervention study confirmed that positive modulation of these pathways could restore proper epithelial-barrier structure in the mouse model, suggesting a potential treatment path in humans given the molecular and histopathological similarities. A reanalysis of the published data with Metascape identified only a partial, tangential match to the author’s original findings with the collapsing function in use, “common bile duct development” (GO:0061009, −log(P)=3.90). With the collapsing feature ignored, a more direct match was identified as “bile secretion” from KEGG (hsa04976, −log(P)=2.258) (**Supporting Data STable 18-1**). However, this match was p-value ranked as pathway 776 out of 982 in the complete pathway list, likely limiting its human discoverability. No relevant matches to either the NAD+ or bile acid pathways were recognized with ToppGene (**Supporting Data STable 18-2**). GeneAgent reported “regulation of cellular stress and metabolic equilibrium” as the final process summary (**Supporting Data STable 19-3**). In this case, it is highly unlikely that any of the other tools would have enabled the primary hypothesis and subsequent laboratory validation work.

## Discussion

### Novel aspects and capabilities of the system

MIRaS represents a new approach to artificial intelligence systems. Instead of building models for response from vast amounts of information, MIRaS leverages a highly efficient computable memory system (GeMM) to store key components of that information and can draw upon, at will, to infer or “reason” answers to system inquiries in near real time. Utilizing MIRaS, CompBio is able to provide responses to complex multi-omic analysis queries that are not based on human-curated pathway knowledgebases but can do so within statistical confines. To the authors’ knowledge, this combination of capabilities has not been achieved by either traditional pathway tools or more recent AI/ML-model based systems. Furthermore, by design, the system is not hallucination capable, and all results are explicitly traceable to source knowledge. While RAG (retrieval-augmented generation) and CAG (context-augmented generation) systems can provide some degree of traceability, it is typically window limited. As many of the MIRaS capabilities are complementary to limitations of LLMs and other foundational models, a major ongoing development effort in the laboratory is an attempt to integrate agentic AI with the MIRaS memory system to directly leverage the strengths of both.

### Evaluation of the system

Three tiers of system evaluation were provided to demonstrate the capabilities of CompBio and its underlying intelligence engine. Positive control assessments provide a clear illustration of the platform’s ability to accurately identify known biological pathways and processes from knowledgebase lists and experimental data. Comparison with traditional pathway tools on published experimental data demonstrate the platform’s ability to capture the key biological processes with near-optimal efficiency to aid in human interpretation. Finally, in the most critical evaluation, the somewhat unique nature of CompBio’s initial launch and utilization have provided numerous published examples of the system’s ability to drive hypothesis generation with prospective validation.

### System requirements

Detailed system requirements for building the GeMM within MIRaS and running the CompBio application are provided in the Methods section and are quite modest compared to LLM and other foundation model requirements. The GeMM for CompBio is built in less than 24 hours on a single server with a cost of roughly $50,000. An instance of CompBio and MIRaS supporting approximately 400 users operates on an even smaller server costing roughly one fifth the GeMM-building server and neither requires GPUs. Given the potential extensibility and distributability of the MIRaS engine to applications well beyond CompBio, it may provide a complementary, environmentally sustainable path to solutions for many tasks currently being tackled by primarily model-based AI with substantial GPU requirements in large data centers.

### Limitations of the system and future development

While CompBio and MIRaS have already aided in advancing biological research, there are some limitations to the current system. Perhaps the most significant of these limitations is the need to disambiguate short, typically two- or three-character entity names, that are identical to common words or acronyms (e.g. MR – Mineralocorticoid Receptor; MR – Magnetic Resonance). The MIRaS engine currently leverages a fit-for-purpose algorithm to ensure appropriate identification in these cases. While not a limitation per se, the system also requires human provided tables, with associated tokens, to form the knowledge framework dimensions at the initial step of GeMM creation and, in the current generation of the tool, the framework dimensions are capable of utilizing single-token only coordinates, no complex/multi-token (n-gram like) coordinates which, for example, limits the number of metabolites the system currently recognizes. Finally, while the theme generation in CompBio has no dependence on pathway knowledgebases, the labels used to annotate those themes for human readability and interpretability come from, and are limited by, what exists in current biomedical ontologies. Each of these limitations is being directly addressed in the next generation of technology already in development.

## Supporting information

SupportingData_Table1-19

## Acknowledgements

We thank Dr. R. Mitra for detailed, incisive feedback on manuscript revision that sharpened rationale, claims, and scope; Dr. TC Liu for his continued guidance on CompBio development and its application to human disease; Dr. C. Ho for ongoing feedback on algorithm development and manuscript review; A. Wischmeyer for critical technical review of the manuscript; S. Santhanam for project coordination and management; and Dr. J. Gordon and his laboratory for early adoption, implementation, and continued feedback that has had a major impact on the development and visibility of this system.

## Author contributions

RDH, CES and RAB designed the project. WDH, CM, RDH, MRJ, DK, BKS and JML developed the application and coded the algorithms. RAB, CES, RDH analyzed and interpreted the data. JFM generated the figures. RDH, RAB, CES, JML, WDH, CM, DJK drafted the manuscript. RDH supervised the project. RDH, RAB, CES, JML, WDH, CM, DJK, JFM, BKS, and MRJ revised the manuscript. All authors contributed to the writing of the manuscript and approved the submission.

## System Availability

The CompBio system can be freely accessed by non-commercial users at https://gtac-compbio-ex.wustl.edu.

## Methods

### MIRaS construction

The MIRaS engine utilized by the CompBio platform consists of two major components, a **Ge**neralized **M**emory **M**odel (**GeMM**), which contains tokenized episodic and complex memory(ies) constructed from ingested knowledge sources and a **M**emory-based **I**nference and **R**easoning (**MIR**) system capable of pseudo-reasoning from these memories in response to user inquiries. While not yet true, complex, human reasoning, the MIR is a computational system capable of simplistic reasoning based on a highly generalizable method for signal detection from a broad array of data or information contained within the GeMM.

### GeMM definition – knowledge source selection

One of the primary functions of the GeMM is to maintain explicit episodic memory of data or information contained within source knowledge, with context, in an efficient and computationally amenable framework. The first step in the creation of the GeMM (**Supplemental Figure 1**) is the selection of source knowledge for ingestion and the associated vocabulary(ies) for subsequent computation. CompBio utilizes a GeMM built primarily from PubMed articles and abstracts in conjunction with the vocabularies of biology and medicine. The version 2.9 GeMM is constructed from the Q1, 2025 PubMed release with 38,355,642 abstracts and 6,600,121 full-text articles.

### GeMM definition – episodic-memory framework (EMF)

As knowledge source compendiums may differ extensively in both form and content, a framework capable of storage, maintenance, and computation on the data must be adaptable and generalizable for non-bespoke utilization. To achieve this, an episodic-memory framework (EMF) is defined as a superset of framework dimensions wherein each dimension consists of a set of related tokens that have peer status (e.g. biological entities - genes, metabolites, drugs). In turn, each token represents a specific coordinate, or index, within a given framework dimension. As a note, the framework coordinate system maintains distinct differences from cartesian or linear coordinate systems. The positions of two coordinates within a given dimension do not define a distance, simply two unique locations within that dimension. Furthermore, it is possible for episodic memories to exist at multiple coordinates simultaneously within a given dimension. Thus, the EMF is defined as:

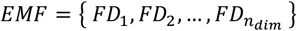

where,

*n*_*dim*_ *=* token sets required to represent source knowledge of vocabulary, ***V***.

Within the EMF, each token set *FD*_*x*_, is an assembly of peer tokens that define the coordinates of a given dimension:

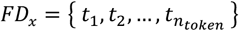

where,

*t*_*k*_ = unique peer-related token coordinate within the dimension.

*n*_*token*_ = number of tokens in *FD*_*x*_, typically derived from a trusted source.

Thus, the component of the target vocabulary(ies), ***V***, specifically represented and understood within the EMF is:

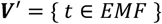

As *V*^′^ → ***V***, the representation of all possible knowledge associated with the given vocabulary becomes more complete but at the cost of system resources that will be described later. The specific framework constructed for the CompBio GeMM is defined as:

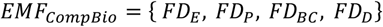

where,

*FD*_*E*_ = Biological Entities (genes/proteins, metabolites, microbes, …)

*FD*_*P*_ = Physiology

*FD*_*BC*_ = Biological Concepts

*FD*_*D*_ = Drugs/Compounds

The biological entity dimension is further granularized to peer subtypes of genes/proteins, metabolites, microbes, and miRNA with genes/proteins and metabolites also having recognized synonyms. The trusted sources utilized to create the CompBio framework dimensions are provided in **Supplemental Figure 1**. Though not leveraged in CompBio, other GeMMs have been constructed with dimensions for knowledge sources such as medical records including, but not limited to, temporal (date based), diagnosis (ICD10), and procedural (CPT) vocabulary components.

### GeMM definition – knowledge sources

With a complete EMF definition, the system can ingest relevant knowledge sources. We define an unfiltered knowledge source, *ks*_*UF*_, as a set of unfiltered tokens:

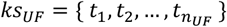

For the ingestion process, we apply a collection of token filters or characteristic functions, *f*_*k*_ (*t*) ∈ {*True, False*}, defined as:

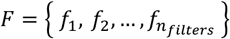

Where we use the notation *f*_*k*_ as shorthand for *f*_*k*_(*t*). Application of these filters on an unfiltered knowledge source, *ks*_*UF*_, yields a filtered token set, *ks*_*F*_ :

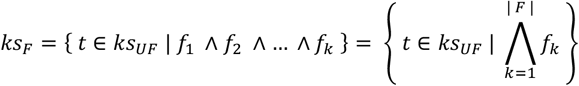

A specific characteristic to the creation of the CompBio GeMM is that a given knowledge source must contain at least one token from the biological entity dimension for ingestion. For PubMed sources, an abstract creates a single source (episodic) memory while full-text articles can, typically, generate dozens of source memories based on a fixed-length text window encompassing each biological entity instance.

### GeMM generation – episodic memory formation

For efficient storage and computation, each filtered knowledge source *ks*_*F,y*_ is converted into an episodic memory, *em*_*y*_, within the knowledge framework:

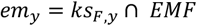

Each episodic memory *em*_*y*_ is further vectorized within the definition of the *EMF* dimensions, 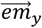, by applying a feature map, *ϕ*_*W*_(*ks*_*F,y*_), that weights each token *t*.

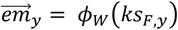

Specifically, within the CompBio GeMM, the token vectors for an abstract receive an initial weight of 1.0, results sections receive a weight of 0.25 and discussion receives a weight of 0.1. Introduction and Methods sections are not utilized for CompBio. This is done to minimize the effect that referencing prior art can have on data momentum and the emphasis of replication on the discovery of independent observations. The knowledge-source episodic memory is the fundamental unit of encoded knowledge within the GeMM. **Figure 2a** provides an example of abstract ingestion.

### GeMM generation - complex memory formation

Due to their tokenized and vectorized construction, episodic memories can be combined by common or related coordinates to form complex memories. A complex memory 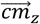 is defined as a combination, or aggregation, of episodic memories that share a common token or logically related set of tokens, *S*, within the *EMF*:

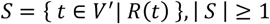

where,

*R*(*t*) = a function that ascribes the relative logical association of token *t*.

Hence, a complex memory is formally defined as:

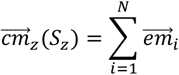

where,

*N* = the number of episodic memories containing *S*_*z*_.

For simplicity, we use the shorthand 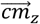 to represent 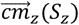. Within the CompBio GeMM, a typical complex memory 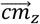 can be an aggregation of a few to 10s of thousands of episodic memories. Subsequently, the set of complex memories associated with the *n* coordinates of a given framework dimension *FD*_*x*_ is:

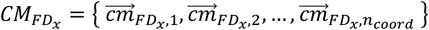

where,

*n*_*coord*_ = number of unique coordinates in framework dimension *FD*_*x*_.

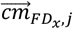 = unique complex memory aggregated on the token at coordinate *c*_*j*_.

Hence, the collection of complex memories across all *EMF* dimensions is:

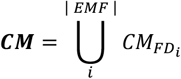

The MIRaS primarily utilizes complex memories, either those computed for dimension-coordinate pairs immediately after source knowledge ingestion or context-specific, to address user-initiated inquiries. Only fit-for-task computations, such as the determination of conditional probabilities for knowledge map generation, described later, occur directly on the episodic memories. A highly simplified illustration of this process is provided in **Figure 2** for the biological concept coordinate of “tight_junction”, 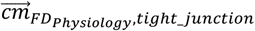.

### GeMM access – memory as a service

The union of all episodic memories 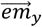 and complex memories 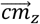 across all framework dimensions define the generalized memory model. Thus, the fully ingested GeMM is:

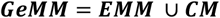

where,

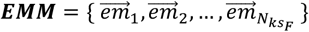

and,

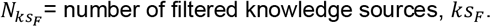

Upon generation, the GeMM is loaded into the active memory of the server and is accessed through an API. As such, the GeMM can be classified as memory-as-a-service.

### MIR Engine definition - memory-based inference and reasoning

Unlike memory-based AI systems that primarily infer based on similarity, the engine within MIRaS also works from differences in memories to identify enriched signal that may be selective or unique. Within this paradigm, the existence, or even magnitude, of a given vector component, which can number in the 10s to 100s of thousands for complex memories, does not directly imply meaningful signal. For example, the majority of complex memories for human genes may be associated with the biological concepts of “microarray” or “RNA-Seq” simply because most have been measured and/or reported within the context of those technological platforms. This is an obvious association with little intrinsic value for discovery. A further complexity is that the determination of what defines “signal” may be contextually dependent and cannot be defined prior to user inquiry. Thus, the identification of signal within memories requires normalization for comparison. Normalized memories are determined by comparing enrichment of a given memory with an associated “peer” memory of the same type and origin. Within the EMF, it is possible to accomplish this for memories of nearly any complexity level and construction. This ability is foundational for the **M**emory-based **I**nference and **R**easoning (MIR) system to respond to complex inquiries.

### MIR Engine–complex memory normalization (semantic memory formation)

Within the *MIR*, memory normalization occurs through comparison of a given complex memory 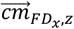 to “peer” memories to identify enriched components selective or unique to that memory. For the set of complex memories associated with each of the coordinates of the EMF dimensions, this is accomplished with the establishment of a memory peer group, *PG*, assembled from randomly selected complex memories residing within the same framework dimensional complex memory set 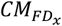:

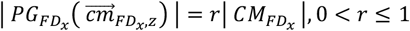

Within the CompBio knowledge engine, 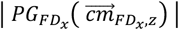 is 40% of all complex memories in 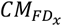 with the exclusion of 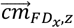 itself. Utilizing the standard formulation for memory aggregation described earlier, all complex memories within the peer group are aggregated into a single complex memory 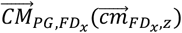. All components of the peer group vector are then divided by the total number of memories aggregated to generate the mean peer group memory:

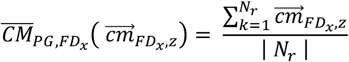

where,

*N*_*r*_ *=* number of randomly selected complex memories in 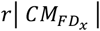.

Illustration of complex memory normalization is provided in **Figure 2c**. Hence, we calculate the normalized complex memory 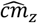, within a given framework dimension, *FD*_*x*_:

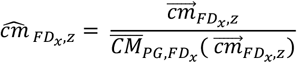

As the formed, normalized complex memories present enriched associations across all underlying episodic memories, they can be considered as semantic memory. This representation allows us to normalize memories for contextual and higher-order complexity as will be discussed next.

### MIR Engine – contextual inquiry

As described, GeMM is capable of storage, pre-computation, and subsequent inquiry responses across a multitude of data and information types based on the constructed knowledge framework. For clarity, the remainder of the description presented here will focus specifically on the knowledge core built to underwrite the CompBio multi-omic analysis platform and will use terminology specific thereto. CompBio is a system designed to aid in the interpretation of high-plex, multi-modal biological data. These data can consist of a single biological entity type, such as RNA-Seq, or multiple entity types such as RNA-Seq and metabolomics. The user inquiry sets are generally derived from experimental data of interest and may represent differences between case and control samples, correlation with phenotypic data, or any of a multitude of experimental analysis approaches. As CompBio does not leverage pre-formed entity lists from curated pathway ontologies, it is not necessary to separate different entity types when processing true multi-omic data (multiple omics entity types from common samples, subjects or conditions). Once a user defined entity set has been selected, it is passed through the UI (described later in the methods) to the knowledge engine as a contextual inquiry. The results of this inquiry will be returned to the user in the form of a knowledge map (clustered knowledge graph) based on information and data previously loaded into GeMM.

The user entity set, *UE*, is defined as:

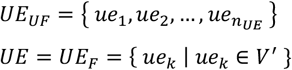

For CompBio, an entity from the user’s inquiry list is deemed valid for inclusion if its associated complex memory that has been formed from three or more episodic memories. Generation of a knowledge map representing the result of user inquiry is a multi-step process that leverages the same complex memory formation and normalization approach described within the knowledge core. First, all user-entity associated complex memories, 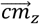, are collected from the knowledge core to form the inquiry-related set of memories. These memories are then aggregated into an inquiry-level complex memory (**Figure 3**).

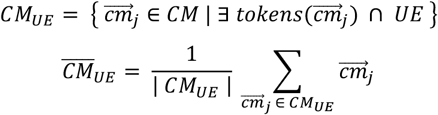

As an inquiry-level complex memory is structurally equivalent to any episodic or complex memory within the knowledge core (simply larger/more complex), the method for identifying enriched, or signal containing, components is also algorithmically equivalent (**Figure 3**). To accomplish this, it is only necessary to define an appropriate peer-entity group to which 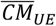 can be compared. To account for the combinatorial space of molecular entities that users can enter in inquiry lists, the system will recognize the specific composition of *UE* at the time of inquiry and build randomly generated sets, or random-peer groups, *RPG*_*x*_, that exactly match the composition of *UE*:

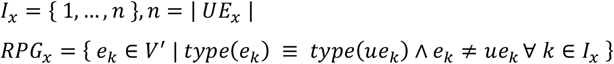

For example, if the user-inquiry list contains 400 genes (RNA-Seq) and 100 metabolites, all random peer groups will contain 400 genes and 100 metabolites randomly selected from *FD*_*E*_.

Assembly of the complex memories on *e*_*k*_ creates the complex memory set for a given random peer group:

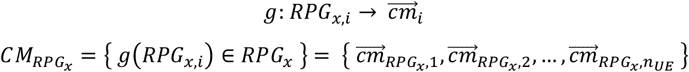

where,

*g* = a function that operates on a single random peer group element, *RPG*_*x,i*_, and can generate an associated complex memory.

For CompBio analyses, 5,000 random peer groups (based on empirical evaluation) are generated for comparison of, and subsequent signal identification within, 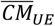. Thus, the complete random peer group complex memory set is:

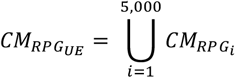

Finally, the mean random peer group complex memory is calculated as:

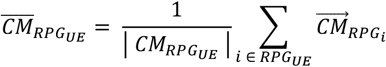

Hence, we can determine the normalized complex memory for the user entity inquiry as:

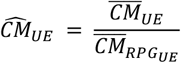

This normalized computation of 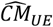 applies the null hypothesis to determine the statistical significance of the entities provided within a user inquiry list against those present within the MIR. Enrichment is calculated for each 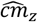 within 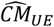:

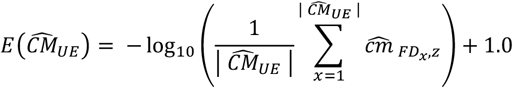

Due to the extensible nature of the MIR, inquiry type and construction are only limited by framework definition and source knowledge ingested within the GeMM. As examples, the MIRaS system has been successfully tested with GeMMs built from medical claims data to identify components of fraud and waste, as well as data from the FDA Adverse Event Reporting System (FAERS).

Thus, MIRaS can be formally defined as the MIR engine operating on the GeMM core:

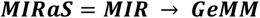

### Knowledge transfer - knowledge map generation

As described, MIRaS is capable of generating both simple and intricate output based on the complexity of the system inquiry. Specifically, in the case of CompBio, the desired output necessitates the creation of a knowledge graph containing descriptors relating to biological processes and how these descriptors are related to the entities in the user inquiry. However, the raw knowledge graph requires a level of abstraction, or translation, to properly convey this knowledge in a human interpretable form. Furthermore, CompBio is designed not only to optimize the simple conveyance of the resulting information but to also enable user learning and interpretation in the form of an explorable knowledge map. To this end, a set of post-processing tools have been constructed enabling computer-human interaction and knowledge transfer.

### Knowledge map - generation

Knowledge map generation occurs in a multistep process that first identifies component interrelationships that can be utilized to form higher order constructs. The purpose is to simplify mapping and visualization of the computer-generated knowledge for human interpretation. The first step in this process generates a conditional probability matrix for all enriched coordinate vectors, henceforth referred to as “concepts”, identified by MIRaS. The complete list of enriched concepts (*N*) form a *N* × *N* matrix which stores the mean conditional probability of concept pairs based on presence and association within corresponding memories The matrix conditional probability elements are computed, initially, by assembling a list of all episodic memories associated with each concept revealed in the MIRaS analysis. As the system is predicated on contextual accuracy, conditional probabilities are computed only from the source knowledge utilized in evaluating the user inquiry, not the entirety of GeMM. Subsequently, the mean conditional probability is computed for each concept pair *c*_*i*_ and *c*_*j*_ as:

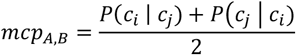

Once computation of all mean conditional probability values has been completed, the concept conditional probability matrix is trimmed, sorted, and trimmed again to enrich the top elements of the matrix for information density (signal and connectivity).

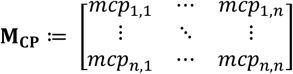

To optimize the efficiency of knowledge representation in the resulting map, the **M**_**CP**_ is first sorted and then trimmed utilizing a factor referred to as the information density score (IDS). The IDS identifies concepts that simultaneously contain the highest levels of signal enrichment and the highest degree of interrelation with other concepts. A bi-directional sort (rows and columns) is then executed on the matrix in descending order with the concept having the highest IDS score at position 1,1. The IDS is calculated as follows:

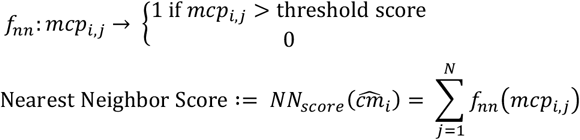

where,

*fnn* = the function that calculate the nearest neighbor score.

Next, we define and apply ranking functions to calculate the IDS:

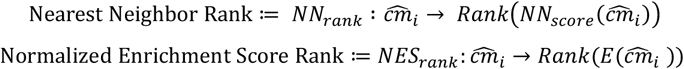

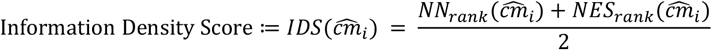

After sorting, the matrix is trimmed to the upper 60th percentile of the concepts. This cutoff was empirically derived for CompBio based on an assessment of thousands of knowledge maps generated from a wide range of sparse to dense input matrices. Concepts that fall below this cutoff tend to form loose themes with little connectivity and are thus removed from further consideration in the resulting knowledge map.

To generate a visual representation of the derived concepts, the sorted and trimmed concept conditional probability matrix is utilized with an accompanying unit matrix of the same rank, *J*, to compute a pseudo-distance matrix defined as:

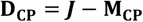

Within CompBio, the resulting distances are used to create biological themes through clustering. The clustering method first uses Isomap^43,44^ (seed independent) embeddings to generate initial 3D coordinates. Concept coordinates are further refined with a force-directed simulation, and final clusters are formed using DBSCAN^45^, a spatial density-aware cluster formation procedure. The result are sets of strongly interrelated biological concepts (biological themes) that were formed from the knowledge sources specific to the user entity list, as opposed to the entirety of the knowledge source corpus, providing contextual awareness. Additionally, the formation of biological themes is further refined through holistic awareness of the biological space yielding a highly efficient representation of the pathways and processes identified, leaving little to no redundancy as is common with ontology-based pathway tools.

3-Dimensional spatial coordinates for resulting biological themes are computed from the centroid of coordinates for concepts within that theme. This is the final step abstracting the low-level knowledge graph into a human digestible and interrogable knowledge map for which the theme coordinates provide the foundation. Based on the criteria provided for theme formation and positioning in the map, it is somewhat self-evident that the distance between themes is a further indication of their degree of relation. As a note, while entities can be associated with any number of biological themes, the concepts from whence the themes are formed can only exist within a single theme. This is a key feature underwriting the limited redundancy observed in CompBio knowledge maps.

Finally, a theme-level enrichment score is computed utilizing the enrichment scores of both the concepts and entities associated with each given theme:

Given the concepts that belong to a theme, *T*_*x*_ :

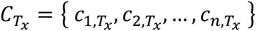

And, the biological entities *BE* associated with *T*_*x*_:

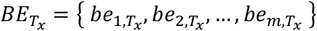

Hence, we can compute the enrichment score *ES* for *T*_*x*_, as:

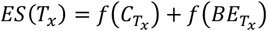

*where*,

*f* = function that aggregates the enrichment score from a theme.

The themes are ranked and sized, within the visualization tool, by the raw enrichment score. The ranking of the theme will be further used in the calculation of statistical significance.

### Knowledge map - labeling

As themes within a CompBio knowledge map are generated *de novo* from source knowledge, without the use of biological pathway or biological process knowledgebases, annotation of these themes would require either the use of LLMs or a sophisticated approach to utilizing existing biological labels from a multitude of trusted sources. While LLMs are powerful utilities, the latter was chosen for two reasons. First, to provide labeling that users are accustomed to from decades of pathway tool utilization and second, to provide individual theme annotation that is contextually aware of the entire knowledge map. To the authors’ knowledge, the latter is a unique feature of CompBio. Approximately 88,000 biological labels from trusted sources HUGO, ENSEMBL, UNIPORT, JAX, mirTarBase, HMDB, and NCBIs MESH^46–50^ have been accumulated to support the theme annotation process. Utilizing a process similar to that described in EMM generation, labels are tokenized and mapped to individual themes and the trimmed **M**_**CP**_ to identify local and global context preservation. A scoring system that includes both the local and global components creates a rank order list of labels for each theme. If multiple non-redundant labels exceed the scoring threshold, up to three labels, above the confidence score of 1.5 (empirically derived), may be used to completely describe the biology represented within that theme. It should be noted that these labels do not represent possible alternate annotations, but a combined description of the biology within the theme. If no high-confidence label can be identified above the minimum scoring threshold, the theme will remain unannotated. Confidence scores are assigned to each label ultimately associated with each theme. Confidence score ranges are based on empirical assessment of thousands of themes/labels across a broad range of biological data. As such, theme annotations often represent the most context-relevant solution(s).

### Knowledge map – statistical significance

Experimental significance of the thematic knowledge map is determined through an empirical method derived from application of the null-hypothesis at the theme-level. This is enabled in two stages. The first stage occurs during the ingestion of a new knowledge source set. Active biological entities from the most recent *EMF*_*CompBio*_ are extracted and are then used to create randomly generated entity lists ranging from 3 to 2,500 entities. To obtain sufficient sampling for the null hypothesis, groups of random lists are built for each input list size. For input lists less than or equal to 100 entities, 200 random lists are generated for each bin size. Above 100 entities, 20 randomized lists are generated for each bin size up to 2,500. Thus, a total of 67,600 random runs is generated across the entire range, and each randomized list is subjected to a CompBio inquiry to generate a complete knowledge map.

To compute both the normalized enrichment score and p-value associated with themes generated from a user inquiry, each theme is subsequently compared to a dynamic window, based on inquiry input size, of randomized data. The window contains a minimum of 2,000 random comparators and is defined such that there are an equal number of samples spanning a ± delta above and below the inquiry size. These 2,000 sample points define a random peer group for determining theme significance in context of the null hypothesis, similar to the previous calculations involving random peers. The enrichment score of each theme from the user-inquiry knowledge map is then compared to the distribution of enrichment scores from its defined randomized comparators of the identical rank (i.e. Theme 1 enrichment score is compared specifically to the randomized data map distribution of Theme 1 enrichment scores. Theme 2 to Theme 2 distributions and so forth.) Based on the comparator size, a minimum p-value of 0.0005 can be computed for each theme. Experiment-level p-values are similarly computed against the randomized data but utilize the IDS of the associated concept matrix **M**_**CP**_. The normalized enrichment score for each theme is also computed utilizing the theme enrichment score data from the randomized sets as follows:

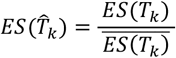

Thus:

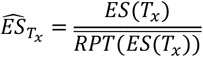

Note that for the randomized entity lists, entity sub-type composition was tested but determined not to impact the score distributions associated with a given inquiry size. Thus, all randomized data lists have been generated using gene/protein entities for simplicity.

### Knowledge map – assertion engine for contextual analysis

Analysis of differential expression patterns is rooted in the comparison of two sets of inputs: a control set (e.g., normal adjacent tissue) and an experimental set (e.g., a tumor biopsy). The signal arising from this pairwise comparison forms the core inner analysis loop of this work, as described in preceding sections. When applying these methods to real-world multi-omics analysis workflows, a scaling problem became apparent, where the scope of contextual comparisons grows (*O*(*n*^2^)) with the number of inputs. Consequently, large analyses were blocked by the problem of accurately matching context across the many possible sets and hierarchies of biology. The assertion generator module inverts the problem to solve it, implementing an outer analysis loop where detailed biological context is automatically matched and used to filter assertions. Thus, full-scale experimental analyses (up to thousands of multi-omic samples) may be productively interrogated together in context. Given a matrix of normalized enrichment scores, the assertion generator automates the construction of contextual assertion graphs. In these graphs, nodes are biological concepts and entities, connected to the network by edges whose length is the contextual distance between them. Densely connected subgraphs occur where there is a shared biological pattern amongst the given inputs. Similarity measures over local and global contexts enable a scientist to test and reject hypotheses at the macro scale. To develop the assertion generator module, we explored many parameterizations, via signal analysis and AI/machine learning techniques (e.g., linear probes over various features, convolutional filters, multinomial logistic regression, neural networks, various ensembles). However, as in many scientific data domains, the difficulty of validating and interpreting results was a key limitation of methods with learned parameters. Observing how the centrality of relevant and biologically conserved subgraphs in a network was the most important factor for explanatory power and biological interpretability, a metric for closeness centrality was implemented. Direct optimization on this metric enables fast, reliable, and explainable results from the assertion generator. To provide a statistical basis for support of a given assertion (versus a null hypothesis), the assertion generator produces p-values and similarity distribution plots. Because the true distribution of biological signals and contexts is unknown, representative and size-matched sets of random inputs are computed. Matching contexts are drawn from the distribution and used to compute similarity scores and p-values. Where the outputs of a given query are clearly distinguishable from noise (e.g., p < 0.01), a spurious match for the asserted biological context is unlikely; where outputs are weakly separated or indistinguishable from noise, there is no basis to support the assertion of a biologically relevant pattern being present in the data.

### CompBio Client – user environment technical description

- **CompBio UI:** Web-based platform developed using Laravel 11 to support computational biology workflows, integrating data analysis, visualization, and research project management. It combines a modern Laravel architecture with legacy PHP components to ensure continuity and backward compatibility during ongoing system modernization.
- **System Architecture:** The application follows a Model-View-Controller (MVC) paradigm, employing Laravel’s Eloquent Object-Relational Mapping (ORM) and Blade templating engine. It is implemented in PHP 8.2+, with SQLite used for local development and PostgreSQL for staging and production. Schema versioning is maintained through Laravel migrations and seeders. Long-running analytical processes, such as the PCMM and Assertion Engine computations, are handled asynchronously via Laravel queues and background PHP worker processes.
- **User and Project Management:** User authentication is implemented using Laravel UI auth scaffolding, with support for lab-based project grouping to facilitate collaboration. Administrative tools allow for metadata editing, project reassignment, and oversight of data access. Notification mechanisms inform users of project completion, system messages and announcements.
- **Frontend Design:** The frontend integrates Laravel Blade templates with JavaScript components. Styling is managed through Bootstrap 5 and Tailwind CSS, using SASS CSS preprocessing. The system utilizes Vite for asset compilation and hot module replacement during development. Axios is used for asynchronous data requests. The transitioning legacy components are utilizing native PHP templates for page content generation, Bootstrap 3 and CSS for styling, and jQuery, jQuery-UI and JavaScript for functionality.
- **Visualization Capabilities:** A central component of CompBio is the interactive visualization module built on Three.JS, a JavaScript 3D Library, enabling users to explore large-scale biological relationships. The visualization pipeline processes raw JSON data through several stages: data normalization, spatial positioning using force-directed algorithms, geometry creation, material application, and final WebGL rendering completion into a Three.JS scene. The visualizer employs multiple renderers layered and synced to a Three.JS Perspective camera and Trackball control module. Point and click interactions are using Three.JS Raycaster that calculates user input in the 3D space, based upon the point-of-view (POV) angle, object to camera distance and scene object render order. Visualization controls and settings are provided by DAT-GUI JavaScript Control Library. Performance optimizations include frustum culling, back-face culling, BufferGeometry, and adaptive layouts for varying screen sizes, resolutions, DPI’s(dots-per-inch). Additionally, various JavaScript code level techniques are being implemented for semi-forced or triggered Garbage Collection within the browser JavaScript engine to minimize memory overhead.
- **Auxiliary Visualization Tools:** The system incorporates additional visualization layers;
  - **CSS3D**: Embeds HTML-based elements in the 3D scene
  - **Google Charts**: Provides supplementary 2D graphing and data charts.
  - **Snap.svg**: Enables generation and editing of high-resolution vector exports
- **Secure AI Integration:** CompBio integrates with a HIPAA-compliant deployment of ChatGPT at Washington University School of Medicine (WUSM), offering AI-assisted analysis within a containerized, data-isolated environment. Prompts and outputs remain confined to WUSM infrastructure.
- **Deployment and Environment Support:** Local development is managed using Laravel Homestead, a Vagrant-based isolated virtual machine environment pre-packaged with Laravel. Production deployments utilize a LAPP (Linux, Apache, PostgreSQL, PHP/Python) stack on Ubuntu Server. The application supports Docker-based containerization and environment configuration via docker-compose. Session management uses a hybrid approach to accommodate both modern Laravel and legacy PHP components. File storage follows a structured hierarchy with defined access controls.

### Hallmark pathways for positive control analysis

To assess the ability of CompBio (v2.9) to identify biological pathways from gene lists, Hallmark gene sets (PMID26771021) were used as input for knowledge map generation. Themes were considered significant if the criteria of p ≤ 0.1 and a normalized enrichment score (NES) ≥ 1.3. Theme annotations and concepts were then compared to the Hallmark pathway names. Matches were classified as perfect, synonymous, or inexact. A perfect match was assigned when the theme annotation included a key term or root of the input pathway name (for example, a theme titled “FXR/PXR/Bile Acid” for HALLMARK_BILE_ACID_METABOLISM). A synonymous match was assigned when no title match occurred, but a matching theme concept was present (for example, a theme titled “PPAR/Lipids/Lipoproteins” containing the concept “adipogenesis” for HALLMARK_ADIPOGENESIS). An inexact match was assigned when no annotation or concept matched the pathway name.

### Signal-to-noise testing with hallmark pathways

To determine the effect of noise on pathway identification, genes from five Hallmark gene sets ranging from 32 to 200 genes (**Supplemental Table 2**) were progressively replaced with randomized noise genes drawn from GeMM 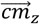 memories, ensuring they corresponded to published genes capable of introducing interference into the pathway signal. Each signal level was tested in triplicate for all pathways. To establish CompBio’s ability to identify these pathways in the presence of noise, serial testing with signal reduction and noise introduction was carried out on five of the original 50 positive control pathways. In each iteration, 10% of the pathway genes were replaced with random genes until 100% of the genes were random and no pathway genes remained. Additionally, three independent replicates were run for each signal to noise ratio iteration as each gene may have a different level of importance to pathway or process that impacts the CompBio theme enrichment score differently (i.e. substituting out a gene weakly associated with the pathway or process will not have the same degree of impact on theme formation as a strongly associated gene).

### Differential gene downstream analysis

Transcriptomic data from drug perturbations of human primary cells were obtained from the GDPx2 functional genomics dataset (https://huggingface.co/ginkgo-datapoints)^23^. Differential expression results (fold changes, p-values, and related statistics) were used as provided. Perturbation-response profiling was performed using DRUG-seq, a 3^′^ digital RNA-seq method with RNA barcoding to reduce amplification bias. Primary human aortic smooth muscle cells and dermal fibroblasts were treated with compounds from the LOPAC1280 library. Two drug concentrations were considered for differential gene selection. The low concentration was 28.5 nM, or the lowest dose that produced a significant CompBio knowledge map (wortmannin low dose was 300 nM). A compound dosage of 900 nM was used for the high concentration. Differential gene selection applied thresholds of fold change ≥ 1.5 in the indicated direction and a nominal p ≤ 0.05. Gene lists were capped to the top 1,500 genes ranked by significance and fold change. These lists were used as input to CompBio, and interpretation was limited to themes meeting significance criteria described above.

### Pathway tools and published datasets

To assess CompBio performance with real world datasets, we used CompBio and three other popular methods Metascape, ToppGene and GSEA with three published datasets.

#### Metascape analysis

The web interface for Metascape (http://www.metascape.org) was used with the following parameters-Custom Analysis was performed with the databases selected for enrichment: KEGG Pathways, Gene Ontology, Biocarta, Reactome Gene Sets, Canonical Pathways, Hallmark pathways, WikiPathways and Panther. For pathway and process enrichment, default parameters were used (Min Overlap = 3, p-value cutoff = 0.01, Min Enrichment = 1.5). The complete output file “Enrichment GO/_Final_GO.csv” was used for computing the number of pathways and genes to pathways. The parameter “FirstInGroupByLogP = 1” was used for identifying the collapsed pathways and those with LogP2 < −1.99 were considered as passing significance threshold.

#### ToppGene analysis

The Toppfun feature within the ToppGene Suite (https://toppgene.cchmc.org) was used for pathway enrichment analysis of all the genelists. Input genes were mapped using HGNC symbols and synonyms. The following databases selected for analysis were namely Gene ontology, KEGG, Reactome, WikiPathways, Biocarta and Canonical pathways. For p-value calculations FDR (B&H) correction was used with cutoff of 0.05 to identify pathways with significant enrichment. The parameter “Hit.in.Query.List” from the output file was used to calculate genes to pathway mapping.

#### GSEA analysis

Gene Set Enrichment analysis (https://www.gsea-msigdb.org/gsea/index.jsp) was carried out using the GSEA V4.4.0 desktop tool with c2.all.v2025.1.Hs.symbols.gmt as the gene sets database with gene set size filters (min = 15, max = 500), number of permutations = 1,000. Gene sets were considered as significant at FDR < 25%.

The input gene lists for each dataset ranged from 122 to 1,056 genes (Mootha et al.: 296, Brass et al. 122; and, Vandussen et al. 1,056) and were generated as follows. For the Mootha et al. dataset, the gene list was generated with nominal p-value < 0.05 and ratio DM:NGT ≤ 0.8 (Diabetes: normal glucose tolerance). For Brass et al. and Vandussen et al., gene lists were supplied in the manuscript material. All the gene lists used here are available in supplementary material (**Supplemental Table 6**). For GSEA, the full dataset input files for Mootha et al. were made available here (https://www.gsea-msigdb.org/gsea/datasets.jsp) and were used as-is.

#### GeneAgent analysis

Gene Agent analysis was carried out using the following interface https://www.ncbi.nlm.nih.gov/CBBresearch/Lu/Demo/GeneAgent/geneagent.html; V1.0:20250418. The output was a text format with the process listed and evidence for each gene contributing to that process. Results are available in supplementary material (**Supporting Data STable 19**).

### System Requirements

The MIRaS GeMM built for CompBio version 2.9 ingested over 38 million abstracts and 6.6 million full-text articles from PubMed into the episodic knowledge framework. Creation of the associated framework-dimension complex memories took approximately 20 hours and 512 GB of RAM on a server with two AMD EPYC 9534 CPUs (128 cores total) and 3 TB of RAM. The resulting GeMM utilizes approximately 70 – 100GB of RAM on the system. Typical system inquiries require approximately 30 – 40 GB of system memory per inquiry (5 to 2,500 input entities) and execute in approximately 2 minutes, on average.

## Figure Legends

**Supplementary Figure1.**
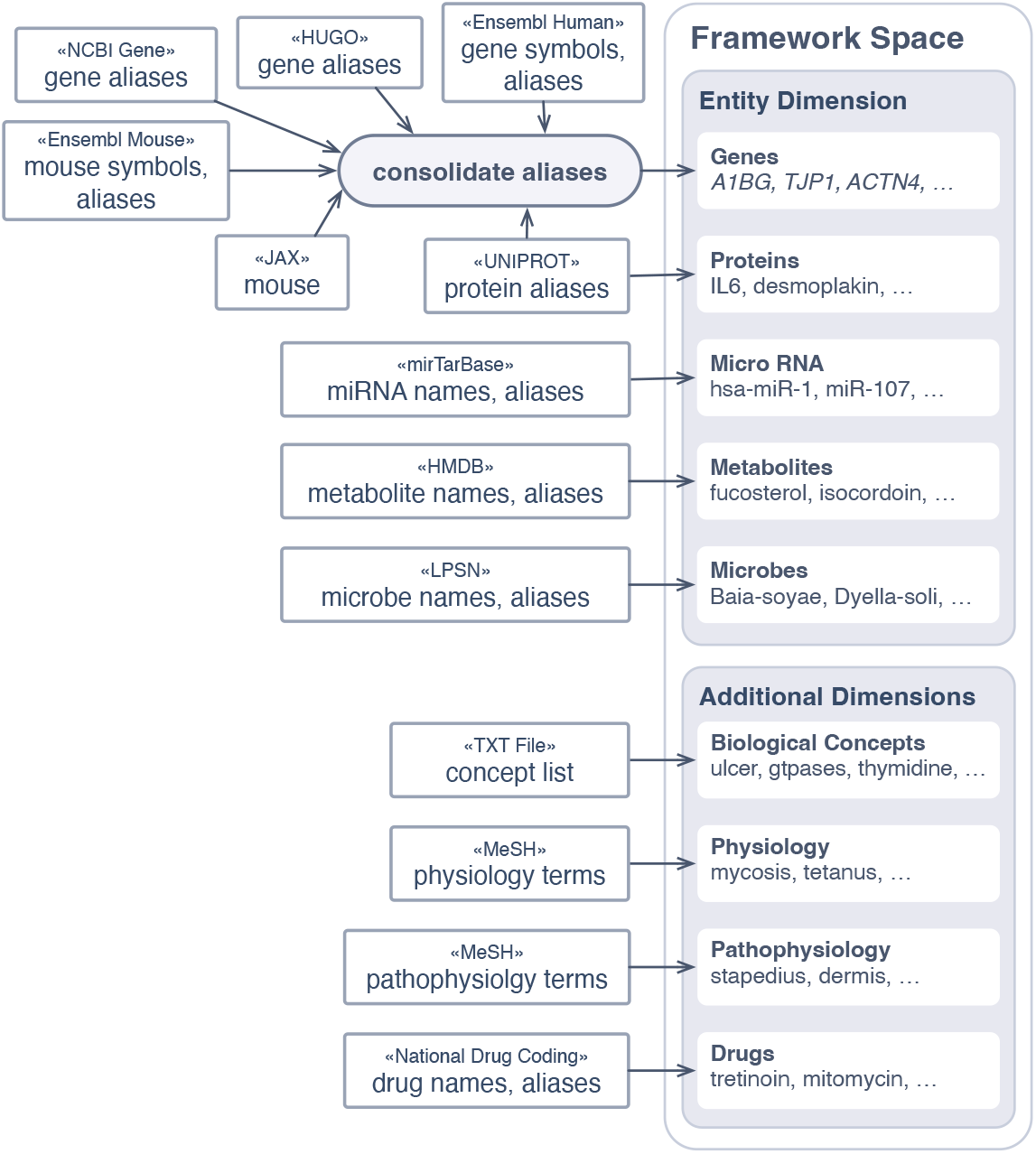
Schematic diagram showing the trusted sources used for generating the CompBIo framework dimensions. The databases and associated fields used and consolidation of symbols is shown.

**Supplemental Figure 2.**
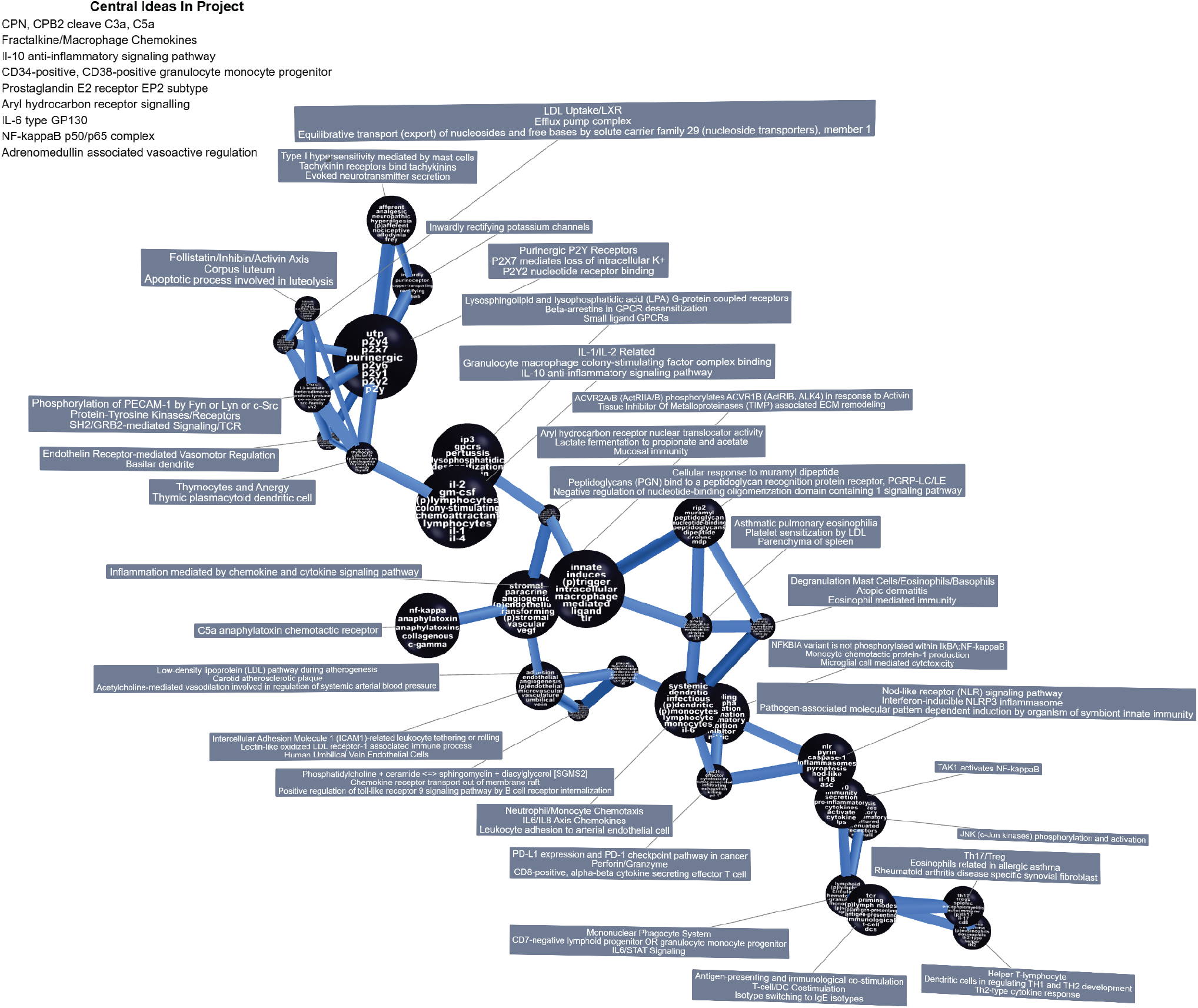
Full CompBio map for the ‘Inflammatory Response’ MSigDB Hallmark gene set. Nodes (“themes”) are clusters of concepts sized by enrichment. Edges weight denotes the degree of shared entities between themes. ‘Central Ideas’ represent major contextually enriched trends within the dataset.

**Supplemental Figure 3.**
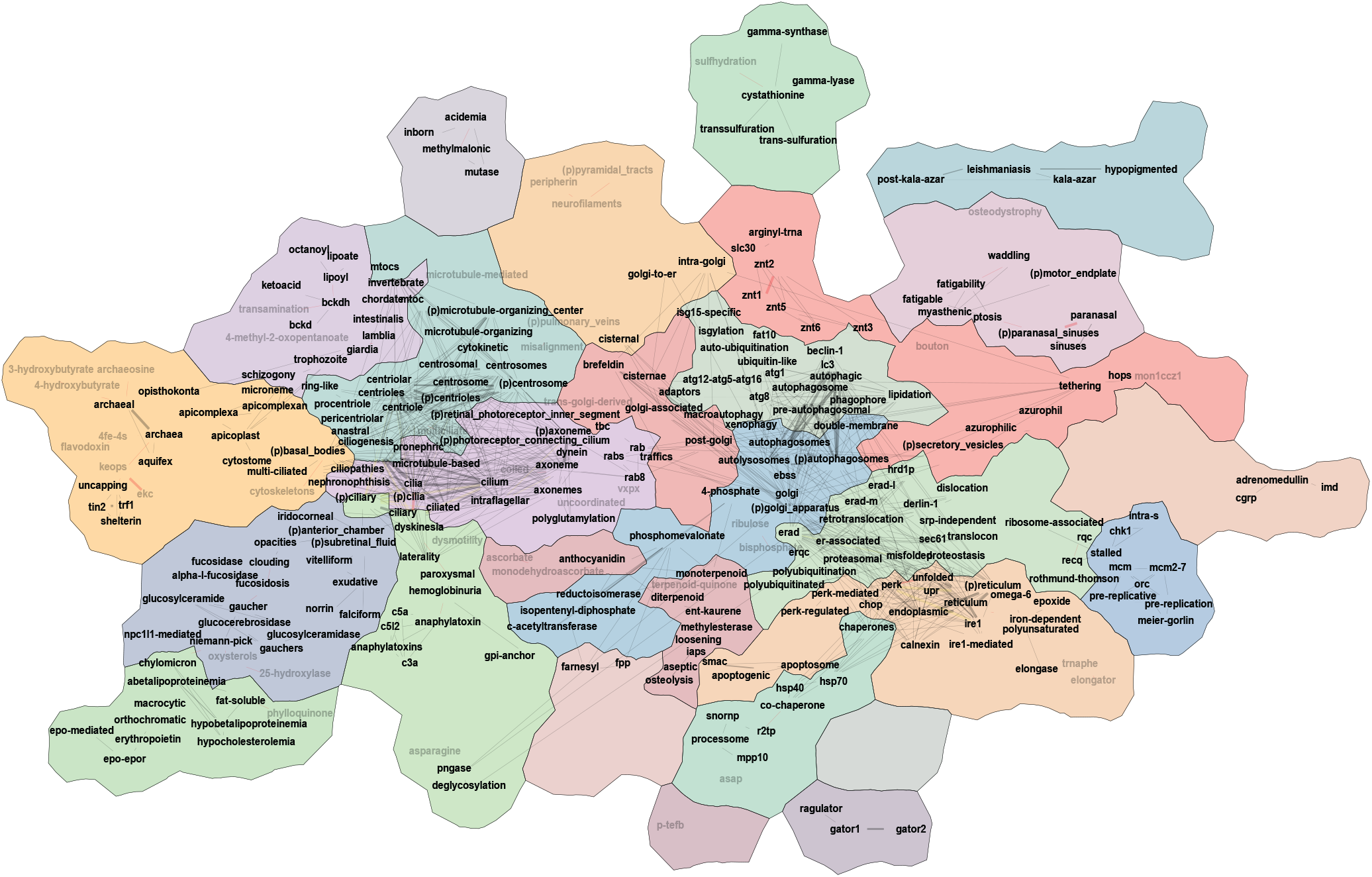
Full assertion engine map comparing the CompBio knowledge maps from aortic smooth muscle cells and dermal fibroblasts treated with 28.5nM brefeldin A. Nodes represent shared biological concepts present in both knowledge maps, and concepts with greater contextual similarity are shown closer together. Edge thickness denotes the degree of cooccurrence of connected concepts in both knowledge maps. Colored ‘countries’ are communities of tightly co-occurring concepts that represent higher-order biological processes.

**Supplemental Figure 4.**
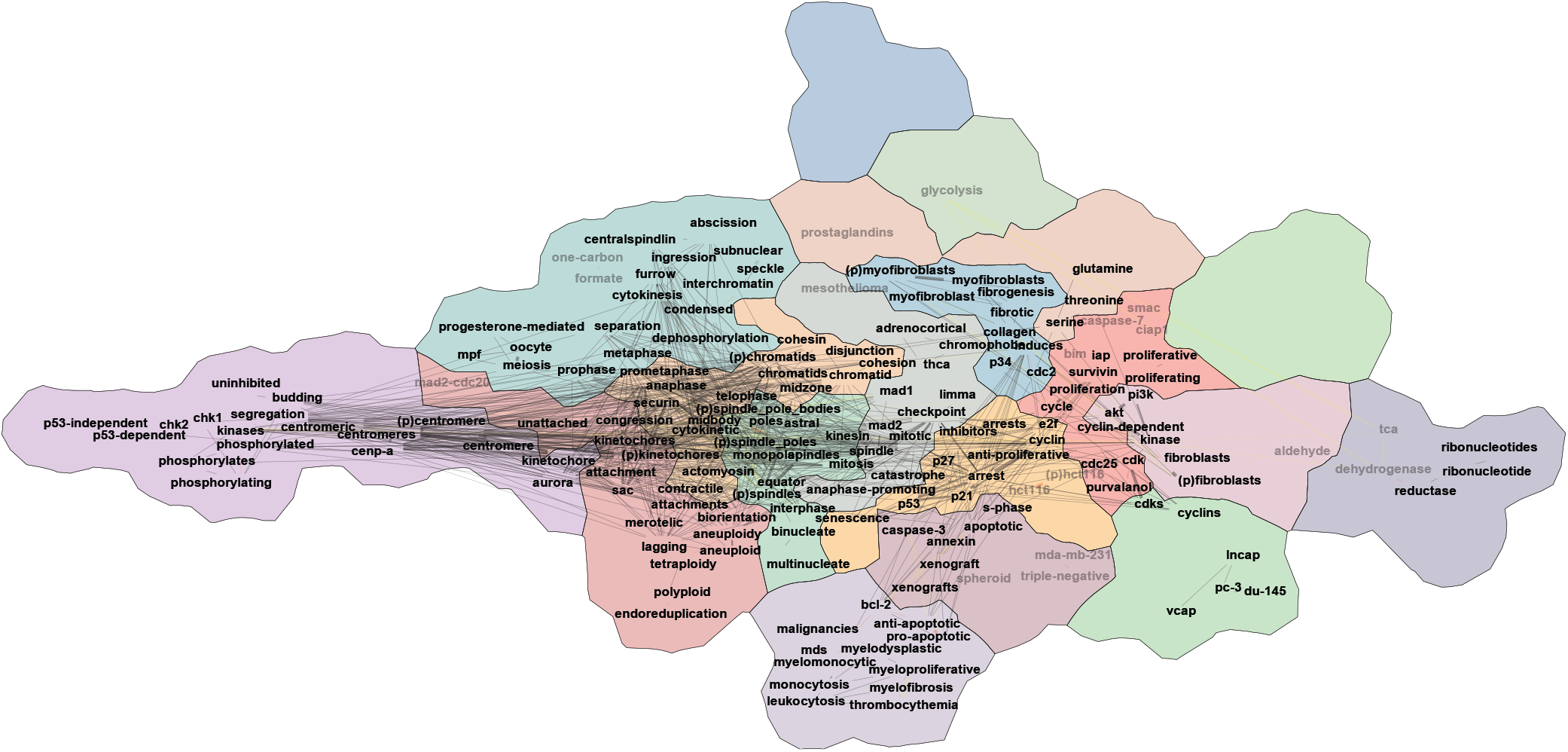
Full assertion engine map comparing the CompBio knowledge maps from dermal fibroblasts treated with 900mM wortmannin or 900mM everolimus. Nodes represent shared biological concepts present in both knowledge maps, and concepts with greater contextual similarity are shown closer together. Edge thickness denotes the degree of cooccurrence of connected concepts in both knowledge maps. Colored ‘countries’ are communities of tightly co-occurring concepts that represent higher-order biological processes.

## References

1 Subramanian, A. et al. Gene set enrichment analysis: a knowledge-based approach for interpreting genome-wide expression profiles. Proc Natl Acad Sci U S A 102, 15545–15550 (2005). 10.1073/pnas.0506580102

2 Mootha, V. K. et al. PGC-1alpha-responsive genes involved in oxidative phosphorylation are coordinately downregulated in human diabetes. Nat Genet 34, 267–273 (2003). 10.1038/ng1180

3 Chen, J., Aronow, B. J. & Jegga, A. G. Disease candidate gene identification and prioritization using protein interaction networks. BMC Bioinformatics 10, 73 (2009). 10.1186/1471-2105-10-73

4 Chen, J., Bardes, E. E., Aronow, B. J. & Jegga, A. G. ToppGene Suite for gene list enrichment analysis and candidate gene prioritization. Nucleic Acids Res 37, W305–311 (2009). 10.1093/nar/gkp427

5 Chen, J., Xu, H., Aronow, B. J. & Jegga, A. G. Improved human disease candidate gene prioritization using mouse phenotype. BMC Bioinformatics 8, 392 (2007). 10.1186/1471-2105-8-392

6 Chen, E. Y. et al. Enrichr: interactive and collaborative HTML5 gene list enrichment analysis tool. BMC Bioinformatics 14, 128 (2013). 10.1186/1471-2105-14-128

7 Jin, Q., Leaman, R. & Lu, Z. PubMed and beyond: biomedical literature search in the age of artificial intelligence. EBioMedicine 100, 104988 (2024). 10.1016/j.ebiom.2024.104988

8 Ashburner, M. et al. Gene ontology: tool for the unification of biology. The Gene Ontology Consortium. Nat Genet 25, 25–29 (2000). 10.1038/75556

9 Gene Ontology, C. et al. The Gene Ontology knowledgebase in 2023. Genetics 224 (2023). 10.1093/genetics/iyad031

10 Jassal, B. et al. The reactome pathway knowledgebase. Nucleic Acids Res 48, D498–D503 (2020). 10.1093/nar/gkz1031

11 Milacic, M. et al. The Reactome Pathway Knowledgebase 2024. Nucleic Acids Res 52, D672–D678 (2024). 10.1093/nar/gkad1025

12 Kanehisa, M. & Goto, S. KEGG: kyoto encyclopedia of genes and genomes. Nucleic Acids Res 28, 27–30 (2000). 10.1093/nar/28.1.27

13 Zhou, Y. et al. Metascape provides a biologist-oriented resource for the analysis of systems-level datasets. Nat Commun 10, 1523 (2019). 10.1038/s41467-019-09234-6

14 Wang, Z. et al. GeneAgent: self-verification language agent for gene-set analysis using domain databases. Nat Methods 22, 1677–1685 (2025). 10.1038/s41592-025-02748-6

15 Cui, H. et al. scGPT: toward building a foundation model for single-cell multi-omics using generative AI. Nat Methods 21, 1470–1480 (2024). 10.1038/s41592-024-02201-0

16 Yang, X. et al. GeneCompass: deciphering universal gene regulatory mechanisms with a knowledge-informed cross-species foundation model. Cell Res 34, 830–845 (2024). 10.1038/s41422-024-01034-y

17 Zhang, H. et al. mosGraphGPT: a foundation model for multi-omic signaling graphs using generative AI. bioRxiv (2024). 10.1101/2024.08.01.606222

18 Delannoy-Bruno, O. et al. Evaluating microbiome-directed fibre snacks in gnotobiotic mice and humans. Nature 595, 91–95 (2021). 10.1038/s41586-021-03671-4

19 Lovell, J. P. et al. Serum Proteomic Analysis of Peripartum Cardiomyopathy Reveals Distinctive Dysregulation of Inflammatory and Cholesterol Metabolism Pathways. JACC Heart Fail 11, 1231–1242 (2023). 10.1016/j.jchf.2023.05.031

20 Madhu, V. et al. The loss of OPA1 accelerates intervertebral disc degeneration and osteoarthritis in aged mice. Nat Commun 16, 5996 (2025). 10.1038/s41467-025-60933-9

21 Sao, K. & Risbud, M. V. SDC4 drives fibrotic remodeling of the intervertebral disc under altered spinal loading. Cell Death Dis 16, 678 (2025). 10.1038/s41419-025-08002-3

22 Valentine, M. C. et al. Multiomic Characterization of Pre- and Post-Neoadjuvant Chemotherapy-Treated Ovarian Cancer Reveals Mediators of Tumorigenesis and Chemotherapy Response. Cancer Res 85, 3558–3570 (2025). 10.1158/0008-5472.CAN-24-3804

23 Winkler, E. S. et al. Human neutralizing antibodies against SARS-CoV-2 require intact Fc effector functions for optimal therapeutic protection. Cell 184, 1804–1820 e1816 (2021). 10.1016/j.cell.2021.02.026

24 Liberzon, A. et al. The Molecular Signatures Database (MSigDB) hallmark gene set collection. Cell Syst 1, 417–425 (2015). 10.1016/j.cels.2015.12.004

25 Baugh, L. et al. Mapping the Transcriptional Landscape of Drug Responses in Primary Human Cells Using High-Throughput DRUG-seq. bioRxiv, 2025.2006.2003.657593 (2025). 10.1101/2025.06.03.657593

26 Lee, U. et al. The Fungal Metabolite Brefeldin A Inhibits Dvl2-Plk1-Dependent Primary Cilium Disassembly. Mol Cells 40, 401–409 (2017). 10.14348/molcells.2017.0032

27 Stilling, S., Kalliakoudas, T., Benninghoven-Frey, H., Inoue, T. & Falkenburger, B. H. PIP(2) determines length and stability of primary cilia by balancing membrane turnovers. Commun Biol 5, 93 (2022). 10.1038/s42003-022-03028-1

28 Wymann, M. P. & Pirola, L. Structure and function of phosphoinositide 3-kinases. Biochim Biophys Acta 1436, 127–150 (1998). 10.1016/s0005-2760(98)00139-8

29 O’Donnell, A. et al. Phase I pharmacokinetic and pharmacodynamic study of the oral mammalian target of rapamycin inhibitor everolimus in patients with advanced solid tumors. J Clin Oncol 26, 1588–1595 (2008). 10.1200/JCO.2007.14.0988

30 Schuler, W. et al. SDZ RAD, a new rapamycin derivative: pharmacological properties in vitro and in vivo. Transplantation 64, 36–42 (1997). 10.1097/00007890-199707150-00008

31 Brass, A. L. et al. The IFITM proteins mediate cellular resistance to influenza A H1N1 virus, West Nile virus, and dengue virus. Cell 139, 1243–1254 (2009). 10.1016/j.cell.2009.12.017

32 VanDussen, K. L. et al. Abnormal Small Intestinal Epithelial Microvilli in Patients With Crohn’s Disease. Gastroenterology 155, 815–828 (2018). 10.1053/j.gastro.2018.05.028

33 Ochoa, J. P. et al. Biallelic Loss of Function Variants in Myocardial Zonula Adherens Protein Gene (MYZAP) Cause a Severe Recessive Form of Dilated Cardiomyopathy. Circ Heart Fail 17, e011226 (2024). 10.1161/CIRCHEARTFAILURE.123.011226

34 Rickelt, S. et al. Protein myozap--a late addition to the molecular ensembles of various kinds of adherens junctions. Cell Tissue Res 346, 347–359 (2011). 10.1007/s00441-011-1281-8

35 Tsukita, S. et al. Roles of ZO-1 and ZO-2 in establishment of the belt-like adherens and tight junctions with paracellular permselective barrier function. Ann N Y Acad Sci 1165, 44–52 (2009). 10.1111/j.1749-6632.2009.04056.x

36 Ungewiss, H. et al. Desmoglein 2 regulates the intestinal epithelial barrier via p38 mitogen-activated protein kinase. Sci Rep 7, 6329 (2017). 10.1038/s41598-017-06713-y

37 Adamo, L. et al. Proteomic Signatures of Heart Failure in Relation to Left Ventricular Ejection Fraction. J Am Coll Cardiol 76, 1982–1994 (2020). 10.1016/j.jacc.2020.08.061

38 Liu, T. C. et al. Western diet induces Paneth cell defects through microbiome alterations and farnesoid X receptor and type I interferon activation. Cell Host Microbe 29, 988–1001 e1006 (2021). 10.1016/j.chom.2021.04.004

39 Tsingas, M. et al. Sox9 deletion causes severe intervertebral disc degeneration characterized by apoptosis, matrix remodeling, and compartment-specific transcriptomic changes. Matrix Biol 94, 110–133 (2020). 10.1016/j.matbio.2020.09.003

40 Silver, R. F. et al. Distinct gene expression signatures comparing latent tuberculosis infection with different routes of Bacillus Calmette-Guerin vaccination. Nat Commun 14, 8507 (2023). 10.1038/s41467-023-44136-8

41 Xia, M. et al. Th9 cells provide protective TB immunity. Front Immunol 16, 1581286 (2025). 10.3389/fimmu.2025.1581286

42 Malique, A. et al. NAD(+) precursors and bile acid sequestration treat preclinical refractory environmental enteric dysfunction. Sci Transl Med 16, eabq4145 (2024). 10.1126/scitranslmed.abq4145

43 Tenenbaum, J. B., de Silva, V. & Langford, J. C. A global geometric framework for nonlinear dimensionality reduction. Science 290, 2319–2323 (2000). 10.1126/science.290.5500.2319

44 Pedregosa, F. et al. Scikit-learn: Machine learning in Python. the Journal of machine Learning research 12, 2825–2830 (2011).

45 Ester, M., Kriegel, H.-P., Sander, J. & Xu, X. in kdd. 226–231.

46 Wishart, D. S. et al. HMDB: the Human Metabolome Database. Nucleic Acids Res 35, D521–526 (2007). 10.1093/nar/gkl923

47 Hsu, S. D. et al. miRTarBase: a database curates experimentally validated microRNA-target interactions. Nucleic Acids Res 39, D163–169 (2011). 10.1093/nar/gkq1107

48 UniProt, C. UniProt: the Universal Protein Knowledgebase in 2025. Nucleic Acids Res 53, D609–D617 (2025). 10.1093/nar/gkae1010

49 Dyer, S. C. et al. Ensembl 2025. Nucleic Acids Res 53, D948–D957 (2025). 10.1093/nar/gkae1071

50 Seal, R. L. et al. Genenames.org: the HGNC resources in 2023. Nucleic Acids Res 51, D1003–D1009 (2023). 10.1093/nar/gkac888

